# Glycosylation-Modulated Conformational Diversity in Neurotrophin Receptors

**DOI:** 10.64898/2025.11.29.691319

**Authors:** Alexandros Tsengenes, Christina Athanasiou, Rebecca C. Wade

## Abstract

Glycosylation is a widespread modification of cell-surface receptors, yet its structural impact is often overlooked due to the difficulty of experimentally characterizing glycans. Glycosylation of neurotrophin receptors has been reported to affect their localization and function. To investigate the effects of glycosylation of the extracellular domains (ECDs) of p75, TrkA, and TrkB neurotrophin receptors, we modeled their ECDs in glycosylated and non-glycosylated states and carried out molecular dynamics simulations of monomeric and dimeric forms of the ECDs with and without a neurotrophin bound. The single N-glycan on the p75 ECD provided minimal shielding and had limited interaction with the neurotrophin, although glycan-glycan contacts between the two p75 monomers may influence the stability of the receptor-neurotrophin complex. In contrast, TrkA and TrkB carry multiple N-glycans that shield the ECDs much more and, particularly for TrkB, increase the contact area between the receptor and the neurotrophin. The p75 ECD was comparatively rigid, independent of glycosylation state, likely due to its extensive network of disulfide bonds. In contrast, without glycans, the TrkA and TrkB ECDs tended to collapse inwards, sometimes obstructing the neurotrophin binding site. Glycosylation of TrkA and TrkB prevented bending of the ECD into more compact states, and instead promoted extended conformations that better accommodate neurotrophin binding. Overall, the simulations reveal distinct, receptor-specific roles of glycosylation in modulating neurotrophin receptor shielding, flexibility, and conformation, with effects on neurotrophin binding.

**Statement of significance:** Neurotrophins are proteins that promote neuronal survival. Their binding to the extracellular domains of transmembrane receptors leads to signal transduction intracellularly. Glycosylation of some neurotrophin receptors has been reported to affect their localization and function but the mechanism is unclear. We investigate the effects of glycosylation on the structure, dynamics and binding properties of the extracellular domains of three neurotrophin receptors by molecular dynamics simulation. We identify distinct receptor-specific effects of glycosylation. In particular, we find that glycosylation of tyrosine receptor kinases prevents concealment of the neurotrophin binding site due to receptor dynamics and favors a more extended receptor conformation, thereby supporting neutrophin binding and the associated signal transduction.

## Introduction

A common attribute of cell surface receptors is the extracellular presence of covalently bonded oligosaccharides – glycans. The glycosylation sites are post-translationally modified during or after protein translation and glycosylation may play a role in proper protein folding, as well as affecting function and cell partitioning. Glycans can be categorized according to the amino acid to which the glycans are attached, e.g., N-glycans conjugate to the N atom of Asn residues whereas O-glycans conjugate the O atom of Ser and Thr residues.^1^ Due to the difficulty of identifying glycosylation sites and glycan types experimentally, the effects of glycans on protein structure and function are often neglected. Here, we therefore employ computational molecular modeling and simulation to explore the influence of glycosylation on the ECDs of three neurotrophin receptors.

Neurotrophin receptors are single-pass transmembrane glycoprotein receptors with a key role in dendritic growth, synapse formation, neuronal survival and apoptosis. Neurotrophins (NTs), the native ligands of these receptors, are homodimeric proteins that bind the ECDs of the full-length receptors and trigger a change in their structure that leads to receptor activation and intracellular signal transduction.^2^ The ECDs bear glycosylation sites, which are known to impact the function of some of the receptors,^3^ but the structural mechanisms, and their potential influence on neurotrophin binding remain elusive. We focus here on the role of glycosylation in modulating the structure and dynamics of the p75, TrkA and TrkB neurotrophin receptors. The p75 receptor is the non-specific (pan/death) receptor that can bind several types of neurotrophin between its ECDs, which are composed of four cysteine-rich domains with 12 disulphide bonds that form a disulphide ladder. TrkA and TrkB are specific receptor tyrosine kinases, with TrkA binding to nerve growth factor (NGF) and TrkB binding to brain-derived neurotrophic factor (BDNF) or NT-4/5. Their ECD is composed of five domains: domains 1 and 3 are cysteine-rich, domain 2 is leucine-rich, and domains 4 and 5 have an IgG fold. Neurotrophin binding between the D5 domains of the two monomers in the receptor homodimer results in signal propagation through the transmembrane (TM) domain and phosphorylation of the intracellular kinase domain.

The extracellular part of the p75 receptor has both N- and O-glycans, with a single N-glycan close to the N-terminal cysteine-rich domain and several O-glycans (the exact number is not known) on the flexible linker connecting the ECD and the TM domain.^4–11^ In contrast, the Trk receptors have multiple N-glycosylation sites on their ECDs but lack O-glycans.^5,9,12^ There is also evidence of glycosylation of the pro-domain of pro-neurotrophins.^13–15^ The single N-glycosylation of p75 has a complex glycan type,^4^ while its O-glycans in the linker were found to contain sialic acids and have the core: Galβ1- 3GalNAc-O-Ser/Thr.^16^ Whereas O-glycosylation of p75 has been suggested to help with the proper folding of the EC linker^7,16^ and the apical sorting of p75,^6–8^ the role of the N-glycosylation of p75 is not clear, as its deletion does not have an effect on receptor cell partitioning.^6,11^ In contrast, TrkA N-glycosylation is important for receptor localization to the plasma membrane and ligand-dependent activation.^3^ When the receptor is not glycosylated, it is trapped intracellularly and it is constitutively active. Thus, it cannot trigger the Ras/Raf/MAP kinase cascade, which initiates in the plasma membrane, and consequently it cannot promote neuronal survival.^3^

Experimental structural biology methods, such as x-ray crystallography, often require removal of glycosylation from the protein structure during sample preparation to increase resolution, as glycans are very flexible and have a lot of structural variability and therefore tend to decrease the X-ray diffraction quality of the crystals or even hinder crystal formation.^17,18^ This high flexibility renders the structural investigation of the effects of glycans on protein structure challenging. A solution is offered by molecular dynamics (MD) simulations, which enable explicit atomistic modeling of glycans and investigation of the dynamics of glycosylated proteins. Advances in mass spectroscopy have facilitated identification of protein glycosylation sites and the respective glycan types, providing a basis for modelling glycans.^19–22^ Here, we have modeled and performed MD simulations of the glycosylated and non-glycosylated ECDs of the human p75, TrkA and TrkB receptors and compared differences in receptor structure and dynamics due to the presence of glycans. Our results indicate different effects and roles of the N-glycans in these three neurotrophin receptors.

## Methods

### Protein modeling

For the modeling of the complex of the p75 receptor with NT-3, the crystal structure of the complex of the Rattus norvegicus (rat) p75-ECD homodimer and the human NT-3 homodimer was used (PDB ID: 3BUK).^23^ The sequence identity with the human p75-ECD is ca. 92% and thus it was considered a good template. This structure contains the rat p75-ECD residues 32-190 (31-189 in human — corresponding to UniProt entries P07174 and P08138, respectively), an N71S mutation in the rat p75-ECD, which results in the serine residue present in the human sequence, and the human NT-3 residues 143-253. It thus comprises (with a few missing residues) the full length of the structured parts of the p75-ECD and NT-3. This structure was processed using the Schrödinger Suite 2020,^24^ by mutating the residues that were different in rat to the human ones using Maestro. Then protein preparation was performed (bond order assignment, optimization of hydrogen-bond network, restrained energy minimization) using the Protein Preparation Wizard.^24^ As the structure was missing the p75-ECD residues 29-30 and the NT-3 residues 139-142 and 254-257 in each monomer, MODELLER 9.23^25^ was used for modeling them. The resulting model lacked hydrogen atoms, so the protonation states were predicted using PROPKA 3.4.0^26^ via PDB2PQR 3.4.0.^27^

For the modeling of the TrkA receptor with NGF, the crystal structure of the complex of the TrkA full-ECD (residues 33-382) homodimer with the mature NGF homodimer (PDB ID: 2IFG) was used.^12^ The structure of NGF was missing the following residues: Ser 122, Pro 182 – Ser 187, Ala 237 – Ala 241 (residue numbering of the UniProt sequence of human NGF, UniProt code: P01138), which were built with the AutoModel class of MODELLER v.9.23,^28^ using default parameters. The linker consisting of residues Pro 182 – Ser 187 has a conserved conformation in existing crystal structures and therefore it was modeled using the structure of human NGF (PDB ID: 5JZ7^29^) as a template. The ECD of TrkA has Ala 33 – Pro 35 missing (UniProt code of human TrkA-II isoform: P04629-1) which was built with AutoModel.

For the modeling of the TrkB receptor with NT-4/5, the full-ECD (residues 32-383) bound to the mature NT-4/5 was modeled. The crystal structure of the complex of the TrkB immunoglobulin-like domain 5 (D5) homodimer with the mature NT-4/5 homodimer (PDB ID: 1HCF)^30^ was used together with the crystal structure of the complex of TrkA-ECD homodimer with the mature NGF homodimer (PDB ID: 2IFG) for the modeling of the D1-D4 domains. The first three residues of the TrkB-D5 are cloning artefacts and were mutated to the wild-type sequence. The rest of the D1-D4 domain residues were modeled with AutoModel using the PDB structure (2IFG) as a template. The structure of NT-4/5 is missing the following residues: Asn 145 – Gly 150 and Gly 207 – Ala 210 in chain A and Gly 81 – Glu 84, Asn 145, Arg 209 and Ala 210 in chain B (residue numbering of the UniProt sequence of human NT-4/5, UniProt code: P34130), which were modeled with the AutoModel class of MODELLER v.9.23.^28^ Additionally, the linker consisting of residues Asn 145 – Gly 150 was built using the structure of human NT-4/5 from the PDB structure (1B98^31^) as a template.

For each receptor, three different systems were modeled; (1) one with the receptor ECD-homodimer bound to neurotrophin, (2) one with one receptor ECD-monomer bound to neurotrophin, and (3) one with the receptor ECD-monomer alone. All systems were modeled with and without glycans (Figure 1, Table S1).

**Figure 1:**
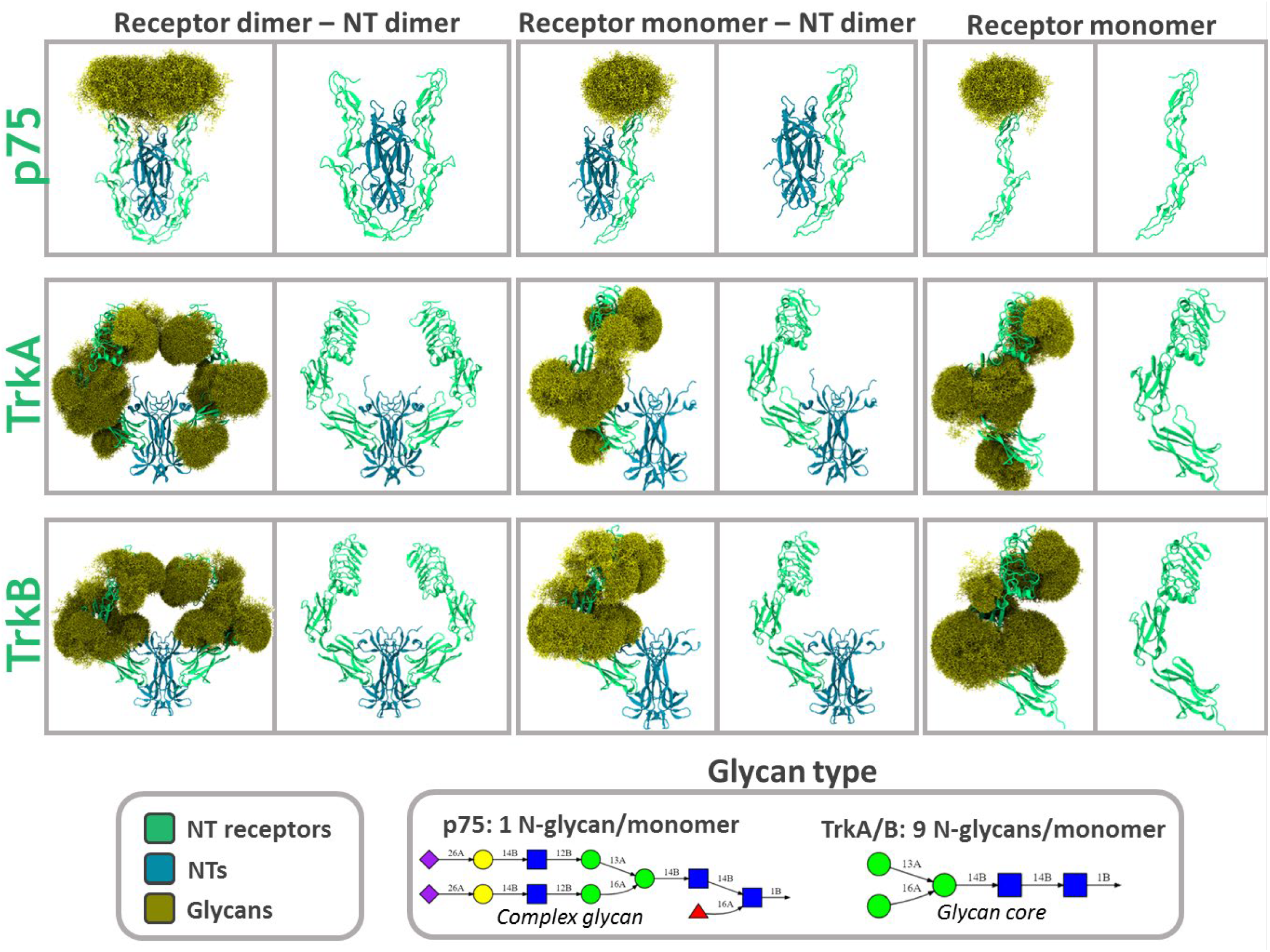
Overview of the modeled glycosylated and non-glycosylated p75 and Trk systems. The systems are: (1) the p75-ECD/TrkA-ECD/TrkB-ECD homodimer with the NT-3/NGF/NT-4/5 homodimer bound, (2) the p75-ECD/TrkA-ECD/TrkB-ECD monomer with the NT-3/NGF/NT-4/5 homodimer bound, and (3) the p75-ECD/TrkA-ECD/TrkB-ECD monomer. All systems were modeled and simulated with and without glycosylation. Proteins are shown in cartoon representation with the receptors in green and the NTs in blue. The systems with glycans present are shown with glycan conformations superimposed from a set of timeframes from the simulations. The glycans are shown in yellow wire representation. The glycan types for p75, TrkA and TrkB are also shown in Symbol Nomenclature for Glycans (SNFG) representation with the blue boxes denoting the N-acetyl-glucosamine, the green circles the mannose, the yellow circles the galactose, the purple rhombus the sialic acid, and the red triangle the fucose monosaccharides.

### Glycosylation modeling

For p75, it is known that each ECD has one complex N-glycan of composition Hex_5_HexNac_4_dHex_1_NeuAc_2_ (Hex: Hexose, HexNac: N-Acetylhexosamine, NeuAc: N-acetylneuraminic acid) (Figure 1) at Asn 60.^4^ p75 also has several O-glycosylation sites, which are located on the linker between the ECD and the transmembrane (TM) domain.^7,8,32^ However, here only the structured ECD of p75 was modeled, so the O-glycosylation was not present.

The TrkA and TrkB receptors have N-linked glycans in their ECD.^3,9^ Specifically, TrkA has 13 potential N-glycosylation sites, based on the Asn-X-Ser/Thr sequon, out of which at least 9 are actually glycosylated.^3^ Six of these sites are known from the crystal structure of the full TrkA ECD bound to NGF.^12^ These are the Asn residues at positions 95, 121, 188, 262, 281 and 358 (Table 3.1). In TrkB, there are 12 potential N-glycosylation sites of which 10 are glycosylated and their exact positions are known; Asn 67, 95, 121, 178, 205, 241, 254, 280, 338, 412 (Table S2).^5,9^ All glycosylation sites in TrkB are in the ECD, except for Asn 412, which is located in the linker between the D5 domain and the TM helix.

To decide on the positions of the glycan sites in TrkA that were not already known experimentally, the possible glycosylation positions in the TrkA structure were compared with the known positions in TrkB. The equivalent positions of the TrkA and TrkB glycosylation sites are given in Table S2. Glycan structures at four additional positions, Asn 67, 202, 253 and 401 (residues in bold font in Table S2), were built in TrkA, resulting in a total of ten glycosylation sites, as for TrkB.

No experimental information is available on the glycan types for either TrkA or TrkB, except for an indication that sialic acid is present in the TrkA glycans.^33,34^ Therefore, we decided to model the glycan root, Man_3_GlcNAc_2_, at all glycosylation sites on TrkA and TrkB that were present in the ECD, i.e. all glycans in Table S2, except for Asn 401 in TrkA and Asn 412 in TrkB, because these are located in the linker after the ECD, which was not modeled. For all systems, the glycans were built using CHARMM-GUI.^35^

### Molecular dynamics (MD) simulations

All glycoprotein systems were inserted into an orthorhombic simulation box with CHARMM-GUI, solvated in TIP3P water with 150 mM NaCl, and then MD simulations were prepared and run using the CHARMM36m force field^36^ and the GROMACS v.2020 MD engine^37^. The protocol consisted of a steepest-descent energy minimization, NVT and NPT equilibration steps with gradual removal of restraints, and one NPT production run. The NVT equilibration step was performed at 310 K for 500 ps with harmonic restraints with a force constant of 400 kJ/(mol·nm^2^) on protein backbone atoms and harmonic restraints with a force constant of 40 kJ/(mol·nm^2^) on the protein side chains. Then, the NPT equilibration was performed at 310 K and 1 atm for 500 ps with restraints on the protein backbone and side chain atoms using force constants of 200 kJ/(mol·nm^2^) and 20 kJ/(mol·nm^2^), respectively. Finally, 3-6 replica production simulations of 500 ns duration were performed for each system without restraints and with different initial velocities for each replica (Table S1).

The Nosé-Hoover thermostat ^38,39^ and the Parrinello-Rahman barostat ^40,41^ were used with a coupling constant of 5 ps and isotropic pressure with 4.5·10^-5^ compressibility. The first equilibration step was run with a 1 fs time step and then a 2 fs time step was used for the second equilibration step and the production run, during which frames were saved every 100 ps. For the treatment of the non-bonded electrostatic interactions, the Particle Mesh Ewald (PME)^42,43^ method was used with a cutoff distance of 1.2 nm, while for the non-bonded van der Waals interactions, a cutoff of 1.2 nm and a switching distance of 1.0 nm were applied. All hydrogen-containing bonds were constrained with the LINCS algorithm.^44^

### Simulation analyses

All simulations were post-processed with the *gmx trjconv* tool of GROMACS to remove periodic boundary condition jumps, reconstruct the molecules and center the proteins in the simulation box. All trajectory frames were aligned to the initial structure using the whole protein backbone. Graphical rendering was done with VMD 1.9.3.^45^ Properties were computed using the MDAnalysis python library^46,47^ and plotted with the Matplotlib python library^48^. The Seaborn python library^49^ was used to compute and plot distributions.

#### Root Mean Square Deviation (RMSD)

The root mean square deviation (RMSD) of the protein backbone was calculated for all systems using the MDAnalysis python library.^46,47^ The RMSD was calculated for all frames of the trajectories and the RMSD time evolution was plotted with the Matplotlib python library.^48^ The distribution densities of the RMSD values were calculated and visualized with the Seaborn python library.^49^ In addition, the internal RMSDs of all protein domains were calculated by aligning the trajectory to the backbone of each domain before the RMSD calculation for that particular domain.

#### Root Mean Square Fluctuation (RMSF)

The root mean square fluctuation (RMSF) of the protein residues was calculated by first aligning the protein Cα atoms to an average structure from the simulations using the align.AverageStructure class of MDAnalysis, which aligns the trajectory to the first frame and then averages the coordinates. Then the RMSF of the Cα atoms of each residue was calculated using MDAnalysis and visualized with Matplotlib. The RMSF was also calculated for each glycan by aligning each ECD protein subunit to the first frame and then calculating the RMSF of each saccharide in the glycans. The RMSF values were plotted in box plots with Seaborn.

#### Radius of gyration

The radius of gyration of the backbone of each receptor subunit in each trajectory frame was calculated with MDAnalysis. The radius of gyration distribution densities for the glycosylated and non-glycosylated systems were calculated and visualized with Seaborn.

#### Solvent accessible surface area

Solvent accessible surface areas (SASA) were computed with the *measure sasa* tool of VMD 1.9.3.^45^ The SASA values of the proteins (both receptor and neurotrophin) were computed every 100 frames with and without taking the glycans into account. The computed values were subtracted to obtain the area shielded by the glycans. Probe radii of 1.4, 2, 3, 4, 5, 6, 7, 8, 9 and 10 Å were used to emulate the interaction with molecules of different sizes, from water to small-molecules and proteins.

#### Contact area between receptors and NTs

The contact area between the p75-ECD and NT-3, as well as the D5 domain of the TrkA and TrkB receptors with the NTs NGF and NT-4/5, respectively, was calculated with a probe radius of 1.4 Å using the *measure sasa* tool of VMD 1.9.3.^45^ Specifically, the solvent accessible surface area (SASA) of the domains of each receptor was calculated twice: with and without the NTs. Then, the two areas were subtracted to retrieve the receptor:NT contact area (Figure S1). For the simulations with the glycans, this calculation was performed with and without the glycans, to identify any participation of the glycans in the contact area. These calculations were repeated for the individual receptor subunits of the receptor dimer systems.

#### Distances and angles between domains

For the receptor homodimers in complex with the NTs, the distances between the centers of geometry (COG) of the domains of the subunits of the receptors were calculated. Specifically, for p75, the D1-D1, D2-D2, D3-D3 and D4-D4 distances were monitored, whereas for the Trks, the D13-D13 (D13 = D1, D2 and D3), D4-D4, D5-D5 and D13-D5 distances were calculated. These distances were compared in the presence and absence of the glycans. Distances were calculated with MDAnalysis and the numpy.linalg.norm function of the Numpy library.^50^ The same python libraries were used to compute the angles defined by the COGs of the D1-NT-D1, D2-NT-D2, D3-NT-D3 and D4-NT-D4 domains for p75 and the D13-NT-D13, D4-NT-D4 and D5-NT-D5 domains for the Trks. Both the distances and the angles were calculated every 100 frames. The data were post-processed with the Pandas library^51,52^ and visualized with violin plots in Seaborn.

#### Principal component analysis (PCA)

Principal component analysis (PCA) was performed to identify the intrinsic hinge motions of the D1 domain of the p75-ECD with respect to the D2-D4 domains, and the D13 domains of the TrkA and TrkB receptors. All the simulation frames were aligned to the backbone of the D2-D4 domains of one p75 subunit or to the backbone of the D4-D5 domains of one Trk subunit. After the alignment, PCA was performed for the coordinates of the D1 of p75 and the D13 of the Trks using the built-in class in MDAnalysis. Plotting of the 1^st^ and 2^nd^ principal component values was performed with Seaborn.

#### Hydrogen bond analysis

The hydrogen bonds (H-bonds) between each glycan and the protein residues were calculated with the HydrogenBondAnalysis class of MDAnalysis for all frames of the trajectories. The results were post-processed with Pandas and Numpy and visualized with Matplotlib as the percentage of the frames in which each glycan forms H-bonds with specific residues. For the plotting, only the glycan-residue pairs which form H-bonds for more than 1% of the frames for p75 and 10% of the frames for the Trks were kept.

#### Dihedral angles (χ1) calculation

The χ1 dihedral angles of the asparagine residues that were glycosylated were calculated for all frames of all replica simulations of both the glycosylated and the unglycosylated systems using MDAnalysis. Plotting of the distributions of the χ1 values was done with box plots in Seaborn.

## Results and Discussion

### Glycosylated models of the ECDs of the p75, TrkA and TrkB NT receptors

The effects of glycosylation of the ECDs of the p75, TrkA and TrkB receptors were studied by modeling and simulating the receptors with and without covalently attached glycans. Different oligomerization states of the systems were modeled: (1) receptor dimers bound to NT dimers, (2) receptor monomers bound to NT dimers and (3) receptor monomers alone, all with and without glycans (Figure 1). Atomistic unbiased MD simulations were then performed for all systems.

The p75 neurotrophin receptor has a single N-glycan of complex glycan type^4^ close to the N-terminal cysteine-rich domain and several O-glycans on the linker connecting the ECD and TM domain.^4–11^ Here, only the single N-glycan of p75 at position N60 of the ECD was modeled (Figure 1).

In contrast to p75, the Trk receptors are known to have multiple N-glycosylation sites on their ECD but to lack O-glycan sites.^5,9,53^ Specifically, TrkA has 13 potential N-glycosylation sites, based on the Asn-X-Ser/Thr sequon, out of which at least 9 are actually glycosylated.^3^ Six of these sites are known from the crystal structure of the TrkA ECD bound to NGF (Table S2).^12^ Of the 12 potential N-glycosylation sites in TrkB, 10 are glycosylated at known positions (Table S2).^5,9^ All glycosylation sites in TrkB are located in the ECD, except for Asn 412 in the linker between the D5 domain and the TM helix. After comparison of the possible glycosylation sites in the TrkA structure with the known equivalent sites in TrkB, four additional glycosylation sites were modelled in TrkA (Table S2). Experimental data on the type of glycans in TrkA and TrkB is lacking, except for an indication that sialic acid is present in TrkA.^33,34^ Therefore, the glycan root, Man_3_GlcNAc_2_, was modeled at all glycan sites on the TrkA and TrkB ECDs (Figure 1).

Visualization of all the glycan conformational ensembles from the simulations reveals some clear differences between p75 and the Trks (Figure 1). For p75, the single complex N-glycan at N60 creates a glycan cloud that shields only the N-terminal part of the p75-EC, where it has minimal interactions with NT-3. On the other hand, the multiple glycosylation sites on the Trks shield several regions of their ECDs and participate in contacts with the NTs. Comparison of the TrkA and TrkB receptors shows that TrkA has glycans over the whole length of its ECD, whereas the glycans in TrkB mostly accumulate closer to its N-terminus. The most important difference is that TrkB does not have any glycan shielding its D5 domain, whereas the glycan at Asn 358 in TrkA covers part of the D5 domain surface. The glycan conformational ensembles do not appear to be significantly influenced by the oligomerization state of the proteins, since no important differences in the glycan clouds are visible between the corresponding receptor monomers and dimers.

The cloud of glycan conformations in p75 appears to block the entrance for NT at the upper region of the receptor, which might affect NT binding and unbinding kinetics. Kinetic studies have shown that NTs dissociate from p75 at different rates, with NGF>NT-3≫BDNF.^54^ Additionally, even though the Trks and p75 interact with NTs with similar dissociation constants in the nanomolar range, the binding kinetics are different, with NGF attaching to and dissociating from the Trks slower than p75.^55^ The differences in the glycosylation profiles between p75 and the Trks could contribute to these kinetic differences.

The root mean square deviation (RMSD) of the protein backbone with respect to the initial structures as a function of time was monitored (Figures S2-S4). For p75, there are no significant differences between the glycosylated and non-glycosylated systems, with only the NT-bound systems exhibiting slightly higher RMSD population values for the glycosylated states (Figure S2). Interestingly, the systems with the p75-ECD as a monomer alone show more fluctuations in the RMSD values, which suggests that the NT might stabilize the p75 ECD. Overall, the RMSD values are below 10 Å, which indicates deviation from the initial crystal structure without large conformational changes.

For the Trks, the RMSD profiles are similar for the glycosylated and non-glycosylated systems (Figures S3, S4), with the “TrkA EC monomer-NT” and “TrkA EC monomer” systems showing slightly higher RMSD values for the non-glycosylated states. In contrast, the “TrkB EC monomer-NT” system displayed smaller RMSD values in its non-glycosylated state, as revealed from the RMSD distribution density peaks. Apart from these small differences, all systems display large RMSD values in the range of 10-30 Å, indicating large deviations from the initial models, which were based on crystal structures.

To examine the origin of these deviations from the starting structures, the RMSD values for the internal motions of each domain of the receptors were calculated, by first aligning to the initial frame of each domain and then calculating the RMSD (Figures S5-S7). The results show much lower RMSD values than the global RMSD calculation - less than 3 Å for most of the domains - revealing that the domain structures are preserved during the simulations. In a few cases, higher RMSD values occurred, such as for the D1 domain of p75, for which the RMSD often exceeded 5 Å (Figure S5), accounting for the higher RMSD values for the full-ECD (Figure S2). Also, the D13-A domain in Replica 1 of the “TrkA EC dimer-NT glycosylated” and Replicas 1, 4, 6 of the “TrkA EC dimer-NT non-glycosylated” systems has high RMSD values. Especially high RMSD values, of ∼10 Å, were observed in Replica 2 of the “TrkA EC monomer-NT glycosylated” system (Figure S6). For the TrkB ECD monomer, whether with or without glycosylation, the D13-A domain mostly had higher RMSD values than the other domains.

A closer look at the sampled conformations of these domains showed that most of the high values originate in the N-terminal linker of p75-D1, which is disordered and quite flexible. Interestingly, the region of the D1 domain where the N-glycan is attached has a more stable conformation, although this also fluctuates (Figure S8). Overall, the rest of the p75-domains have a stable conformation, despite containing several disordered regions. This stability is probably due to the 12 disulfide bonds in the p75 ECD, which form a disulfide bond ladder (Figure S9). The high RMSD values on TrkA occur in the D13 domains, due to changes in the N-terminal region (Figures S10, S11). The D13 domains have several disordered regions, and the N-terminal part undergoes different rearrangements during the simulations. This is in agreement with the high B-factors of this N-terminal region in the crystal structure of the TrkA-ECD bound to NGF (PDB ID: 2IFG)^12^ (Figure S12). Apart from the D13 domains, the D4 domain also sustains conformational changes, especially in the linker between the D and E strands, leading to an RMSD of 10 Å in one replica (Figure S11). Overall, most of the high RMSD values occur for the non-glycosylated systems rather than the glycosylated ones. In general, the RMSD values of the individual domains (D1-D4) are much lower than those of the global RMSD, indicating that the high RMSD values originate in the linker (or hinge) regions between the domains.

Analysis of residue fluctuations during the simulations showed that the glycosylated and non-glycosylated p75, TrkA and TrkB systems behave very similarly (Figures S13, S14) with only the N-terminal region of p75-D1 and the N- and C-terminal residues of the TrkA/TrkB-ECDs showing higher root mean square fluctuation (RMSF) values, since they have more disordered structures. Also, in the “TrkA EC monomer” simulations, the D4 domain on average fluctuates more in its non-glycosylated state (Figure S14C, TrkA). Most notably, the glycans do not appear to influence the intrinsic fluctuations of the Asn conjugation sites as the RMSF values for the glycosylated and non-glycosylated systems are generally the same at these residues. The RMSF values of the NT residues are smaller than those of the receptors, except for the N- and C-terminal regions, which are more mobile because they correspond to unstructured, flexible loops. Also, the glycosylation of the receptors does not seem to affect RMSFs of the NTs over the simulations.

### Differential glycan shielding on p75 vs Trk receptors

The ability of the glycans to bury parts of the receptor surfaces was assessed. As has been previously demonstrated by MD simulations, glycans can shield large parts of the protein area exposed to solvent or other molecules.^19^ Therefore, we calculated the accessible surface area (ASA) of the receptor ECDs with and without glycans, for different probe radii representing molecules of different sizes ranging from 1.4 Å, which corresponds to a water molecule, to 9 Å, which represents a larger molecule, such as a protein (Figures 2A, S15). Intermediate values correspond to the size of small molecules (∼3-4 Å) or oligopeptides (5-7 Å).

**Figure 2:**
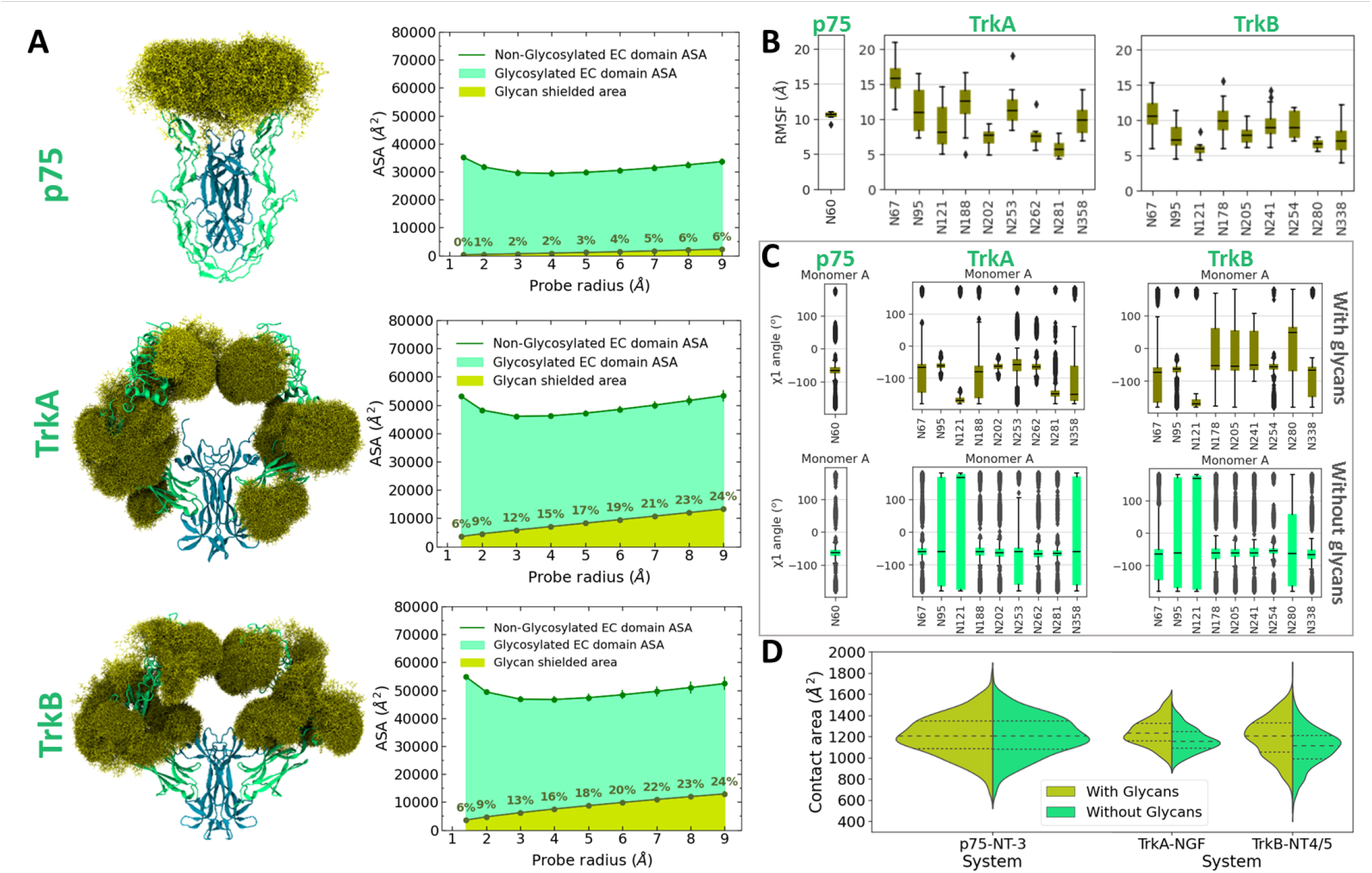
Analysis of shielding and structural changes due to glycosylation for the p75 and Trk “receptor dimer/NT dimer” systems. (A) Accessible surface area (ASA) of the p75, TrkA and TrkB systems (including the ASA of the NTs when present) and the area shielded by glycans computed with multiple probe radii ranging from 1.4 Å (water molecule) to 9 Å (protein-sized molecule). The values were calculated and averaged across all replicas with glycans. The area shielded by the glycans is shown in yellow (with rounded % values), and the green line shows the accessible area of the protein in the absence of glycans. The area highlighted in green remains accessible in the presence of glycans. (B) Distributions of RMSF values for all glycans attached in the Asn (N) residues in the glycosylated p75, TrkA and TrkB systems. (C) Distributions of the χ1 dihedral angle of the glycosylated Asn residues in the glycosylated and non-glycosylated systems. (D) Distributions of the contact area between the receptors and the NTs in the glycosylated and non-glycosylated systems. The quartiles of the distributions are indicated with dashed lines.

Comparison of the glycan shielding among the three receptors showed that the area buried in p75 by the single glycan is much smaller than the area buried in the Trks by glycans at the 9 glycosylation sites, despite the much longer structure of the complex glycan on p75. The reason is that the glycans at N60 on p75 point away from the surface of the protein and tend to interact with each other, potentially contributing to the avidity of the whole complex. Comparison of the TrkA and TrkB receptors shows a similar degree of shielding even though the glycans are attached at different positions on the receptors. This means that the size of the glycans and their total number, which are the same for both receptors, determine the amount of shielding.

This analysis also shows that increasing the size of the probe leads to greater glycan shielding, up to 7% of the receptor ECD ASA for p75 and 28% for the Trks, meaning that the binding of larger proteins may be more disturbed by the glycans than the binding of small molecules. This 4-fold increase in glycan shielding for the Trks compared to p75 might be important for their protection against proteolytic cleavage by e.g. secretases, which has been observed physiologically for p75.^56^

To better understand the origin of the glycan shielding, the intrinsic fluctuations of the glycans were calculated for all receptor systems. For this purpose, each of the domains that have glycans were first aligned to the first frame and then the averaged RMSF values of all monosaccharides of each sugar were calculated (Figures 2B, S16). The glycan on p75 exhibited an average fluctuation of 10 Å with a very narrow distribution, while for the Trks higher RMSF values were observed for the glycans that are located closer to the N-terminus of the ECD. This behavior is in agreement with the lower shielding observed for p75 as higher glycan fluctuations would be expected to effectively bury more receptor surface.

To examine the effects of the glycans on the conformations of the side chains of the conjugated Asn residues, the χ1 dihedral angles of these residues were calculated (Figures 2C, S17-S19). For p75, no significant difference was observed for the N60 residue in the presence or absence of the complex glycan, whereas for the Trks, a more restricted χ1 space was observed when the Asn residues were glycosylated, suggesting that specific conformations of the side chains are adopted upon glycosylation.

Finally, the influence of the glycans on the contact area between the receptors and the NTs was assessed. The contact area was calculated as the difference in the ASA of the ECD with and without the NT present. In the systems where glycans were present, the calculation was performed with and without considering the glycans attached to the D5 domain (Figures 2D, S20). Comparison of the contact area between receptor and NT with and without taking into consideration the presence of glycans shows exactly the same results for p75, suggesting that the N-glycan does not interact with NT-3, whereas for TrkA and especially TrkB, the contact area is higher when considering the glycans, showing that they interact with the NTs. This difference in TrkB is probably due to the glycan attached to Asn 338, which is in close proximity to the NT and which is present in TrkB and not TrkA. Comparison of the ECD dimer with the monomer systems, shows that the contact area per subunit is smaller for the TrkA monomer, while for p75 and TrkB monomers it is similar to that for the dimer (Figure S20). This shows that the packing between NGF and TrkA is tighter in the dimeric system, indicating the importance of receptor dimerization for complex formation with NGF.

### Glycosylation promotes extended conformations of the ECDs of the TrkA and TrkB receptors

The effects of glycosylation on the conformations of the ECDs of the p75, TrkA and TrkB receptors was investigated by calculating the radius of gyration of each receptor subunit. Comparison of the distributions of the radius of gyration of the glycosylated and non-glycosylated systems reveals that the non-glycosylated TrkA and TrkB systems sample conformations of smaller radius of gyration, indicating that the ECD conformations become less extended and more compact, whereas for p75 no difference was observed (Figures 3A, S21-S23). The apparent stability of the p75 conformation suggests greater rigidity of the structure of the p75-ECD, likely due to the presence of the disulfide bond ladder (Figure S9).

**Figure 3:**
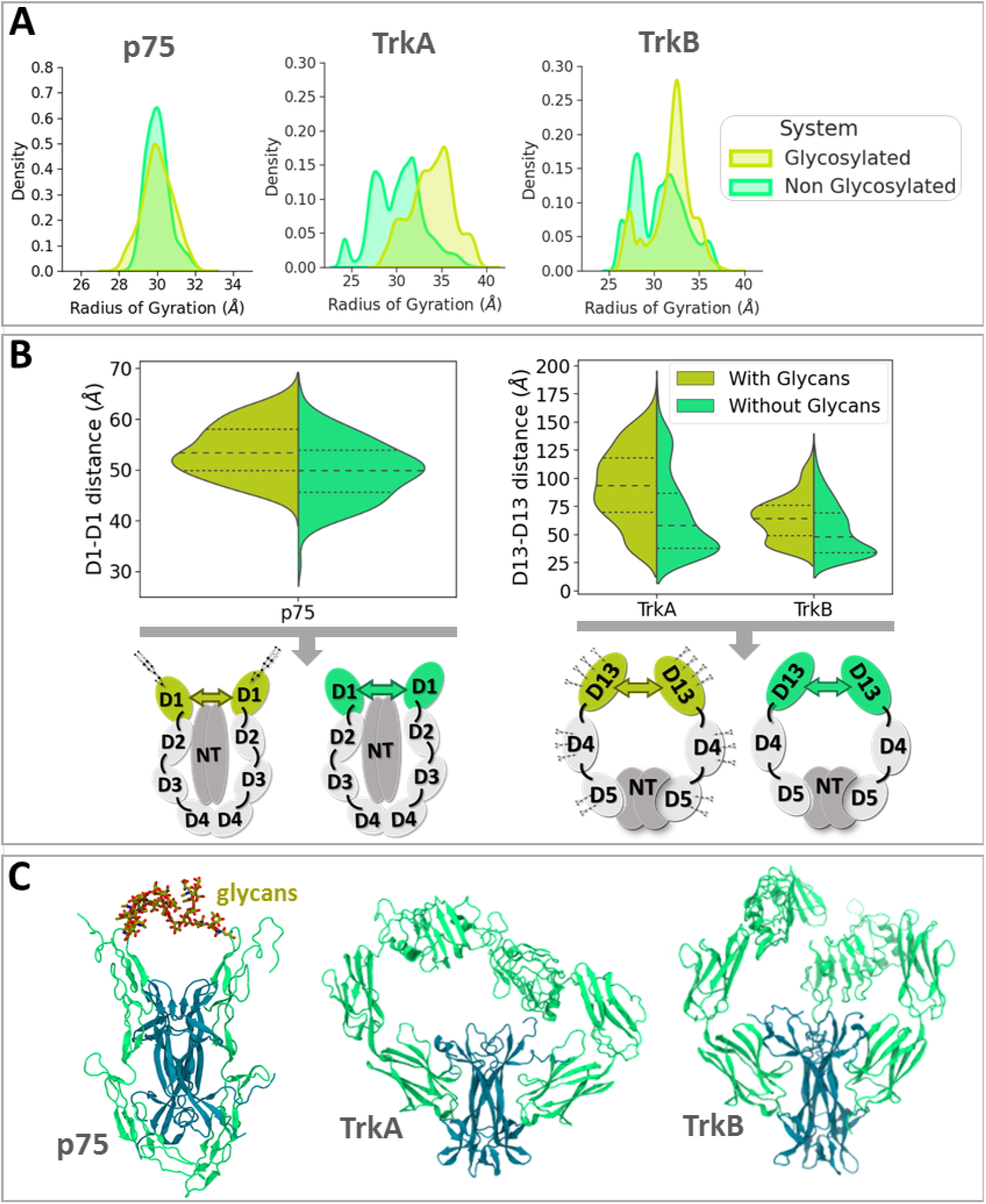
Conformational effects of glycosylation on the p75 and Trk receptors. (A) Distributions of the radius of gyration of the receptor ECD backbone of the glycosylated and non-glycosylated p75, TrkA and TrkB systems. (B) Distributions of the D1-D1 interdomain distances for p75 and the D13-D13 interdomain distances for the Trks. The quartiles of the distributions are indicated with dashed lines. (C) Representative structures showing interactions between the complex glycans at N60 of the two p75-ECD subunits and bent ECD conformations of the non-glycosylated TrkA and TrkB systems.

The extension of the conformations in the presence of glycosylation was more pronounced in the TrkA systems. In the absence of glycosylation, the simulations, which started from the extended crystal structures, showed contraction of the ECDs, allowing sampling of bent conformations. The crystal structure of the ECD of TrkA dimer bound to NGF shows the first three saccharides of the glycans, and thus these may be sufficient to promote the extended conformation of the ECD. Comparison of TrkA with TrkB shows that the effect of glycosylation is less apparent for TrkB. Analysis of the radius of gyration density distributions reveals that glycosylated TrkB samples a broader range of conformations, including those with smaller radii of gyration (< 30 Å), whereas glycosylated TrkA only has conformations with a higher radius of gyration (> 30 Å).

To quantify the hinge motions, the distance between the D1 domains for p75 and the D13 domains for the Trks was computed (Figure 3B). The average values of these interdomain distances are lower for the non-glycosylated systems, with the Trk systems showing a greater difference. The distance distributions between the rest of the domains, as well as the angles among the two domain pairs and the NTs were also calculated, showing that for p75 the D2-D2 and D4-D4 interdomain distances are also a little lower for the non-glycosylated systems, while for TrkA a smaller average D4-D4 interdomain distance was observed in the absence of glycans (Figure S24, S25). The angles follow similar patterns to the distances. These results indicate that the two p75-ECD subunits are somewhat more distant when the two N-glycans at N60 are present, as the two glycans are accommodated between the two p75-ECD subunits (Figure 3C-p75). The glycans in this conformation interact with each other and can potentially block the NT from dissociating from p75, which might affect its binding kinetics. Also, the interacting glycans might enhance the overall avidity of the complex and stabilize it in the homodimeric state. The smaller distances between the D13 domains of the Trks is consistent with the smaller radius of gyration for the non-glycosylated systems and corresponds to bent, closed conformations of the receptors (Figure 3C-TrkA/B, Figure S27). The bent conformations of the Trks were also quantified by the distributions of the D13-D5 interdomain distance (Figure S26), which sample smaller values for all non-glycosylated systems, with this effect being more apparent in TrkA than TrkB.

Principal Component Analysis (PCA) was performed to investigate the main motions of the proteins. For p75, the principal components of the movement of the most mobile domain, the D1 domain of the p75-ECD, with respect to the D2-D4 domains was computed (Figure S28, S29, S30). Comparison of the 1^st^ and 2^nd^ principal components (PC) for the glycosylated and non-glycosylated systems showed similar behavior. The “p75-ECD monomer” system covered more PC1 space than the other systems, which might be due to the increased fluctuations mentioned previously (Figure S13). PCA was also performed on the coordinates of the D13 domains of TrkA and TrkB, as this region showed the largest movements (Figures S31, S32). As for p75, comparison of the principal motions of the glycosylated and non-glycosylated systems shows that they cover mostly the same PC1 and PC2 space.

Finally, an analysis of the hydrogen bonding between the glycans and the protein residues was performed (Figure S37). This analysis shows that, for p75, the Asn 60 glycan makes few hydrogen bonds, most frequently with the adjacent Ala 59. For TrkA, the interaction between the glycan at Asn 202 and Val 200 occurs with > 10% occupancy in all systems and subunits. The glycan at Asn 281 interacts with Gln 279 in all TrkA systems. Indeed, this interaction has ca. 80% occupancy in the second subunit of the TrkA dimer system. Apart from these interactions, the glycans form hydrogen bonds with several Glu, Arg and Gln residues in all TrkA systems. For the TrkB systems, the hydrogen bonds mostly had lower occupancy compared to TrkA. Among the most stable interactions in all systems and subunits was that between the glycan at Asn 205 and Ala 203 of TrkB. The interaction between the glycan at Asn 241 and Trp 230 also occurred with > 20% occupancy. Many glycans interact with Glu and Asp residues on TrkB.

Overall, the analysis of hydrogen bonds between the glycans and the protein residues of the Trks did not reveal any very specific interactions and the glycans mostly interact with their local environment rather than any distant residues. This might be because of the conservative modeling of the glycans with only the first five saccharides (the glycan core). Perhaps the long antennae that are normally present on the glycans under physiological conditions would be able to form more specific interactions. On the other hand, since there is usually a lot of variability in the glycan type according to the cell type, different structures of glycans can be found at a single position. Therefore, one would not expect specific interactions of the receptor with a particular type of glycan structure, as this might not be conserved. Thus, the glycans are most likely to affect the receptor conformations through steric contacts rather than specific hydrogen-bonding interactions.

## Conclusions

The effects of glycosylation on the structure and function of cell membrane receptors are often neglected due to the challenges of characterizing glycosylation experimentally. Here, we modeled the glycosylation on the ECDs of the p75, TrkA and TrkB neurotrophin receptors and performed MD simulations to assess the influence of glycosylation on the receptor structure and dynamics, by comparing glycosylated with non-glycosylated states of the receptors. The three receptors were modeled in three different forms: (1) the receptor ECD homodimer with a neurotrophin (NT) bound, (2) the receptor ECD monomer with a NT bound, and (3) the receptor ECD monomer alone. For p75, it was known that the ECD has a single N-glycan of complex chemical type, which was modeled as such. The TrkA and TrkB receptors, on the other hand, have multiple N-glycosylation sites, which are known for TrkB, and partially known for TrkA. Therefore, we defined the most plausible sites on TrkA by comparison with TrkB. Since the exact glycan types on the TrkA and TrkB receptors are not known, we modeled only the N-glycan core, i.e. the first five saccharides, which are common to all N-glycans.

While glycans are well known to shield protein surfaces, unbiased MD simulations for the glycosylated receptors showed that the extent of shielding differs widely amongst the three NT receptor ECDs. Only the N-terminal part of the p75-ECD could be shielded by the single N-glycan at the Asn 60 position, which also made only minimal interactions with NT-3. TrkA has glycans over the full length of its ECD, whereas the glycans in TrkB mostly accumulate in its N-terminal region. Another important difference is that there is no glycan shielding of the TrkB D5 domain (the domain that interacts with the NT) whereas TrkA has a glycan at residue Asn 358 that shields the solvent exposed surface of the D5 domain.

The ensemble of glycan conformations occupies the inter-subunit space of the upper region of the p75 receptor, and might thereby affect NT binding kinetics, which have been found experimentally to differ for the p75 receptor compared to the Trk receptors.^55^ Glycan shielding was quantified to be minimal for p75 (only up to 7%), while for the Trks it reached ca. 30% of the receptor ECD surface area for large probes, suggesting binding of larger proteins, such as antibodies, rather that solvent or small molecules, might be most affected by the presence of glycans. This may be relevant for protection of the receptors from proteolytic cleavage by e.g. secretases, which occurs physiologically for the p75 receptor.^56^ Another interesting observation was that the glycosylation in the Trk receptors, especially TrkB, formed contacts with the NT, increasing the contact area between receptor and NT, which was not observed for p75.

Investigation of the effects of glycosylation on the conformations of the receptors showed that the glycans promoted the extended conformations of the TrkA and TrkB receptors, while the conformation of the p75-ECD was particularly stable, most probably due to the ladder of 12 disulfide bonds in its structure. Visualization of the TrkA and TrkB trajectories showed that when the glycans were absent, the ECD dimers bend and the two subunits interact with each other or with the NT, whereas in the ECD monomers, the D13 domains interact with the D5 domain. Similar closed conformations of the ECDs have been reported before from enhanced sampling simulations.^57^ Smaller D13-D5 interdomain distances occurred for the non-glycosylated systems, providing direct evidence of the more closed conformations of the non-glycosylated receptors. This was more apparent for the TrkA systems, but also the non-glycosylated TrkB systems seem to explore more conformations with smaller interdomain distances than when glycans are present. For p75, longer D1-D1 interdomain distances occurred in the glycosylated form, likely because the two glycans were accommodated between the two p75-ECD subunits.

Finally, calculations of hydrogen bonds between the complex N-glycan at Asn 60 of p75 and the protein residues revealed interactions mostly with the adjacent Ala 59, as well as Lys 29 and Glu 30. In the homodimeric p75-ECD simulations, the glycans from the two monomers interacted with each other which could stabilize the receptor dimer and contribute to its avidity, and furthermore hinder the NT from dissociating from p75 and affecting its binding kinetics. For the Trks, hydrogen-bond analysis did not reveal any important specific interactions, suggesting that the reason for the glycans promoting the extended conformations of the receptors is mostly due to steric hindrance of the closed conformations when the glycans are present. This is in agreement with the fact that many of the glycans are attached at the linkers of the Trks between the subdomains or close to them. Thus, the glycans can limit the hinge motions of the receptors and keep them in more extended conformations, which might facilitate NT binding. In the systems where the NT and glycans were not present, the D13 domains were able to bend and interact with the D5 domain, which is the binding site of the NT. This bending could hinder NT binding under physiological conditions or reduce the rate of binding and thus receptor activation by the NT.

Previously, we observed in MD simulations that differences in the sequences of the TM domains of p75, TrkA and TrkB result in distinct signalling mechanisms through the TM helix homodimers^58^. Here, this MD simulation study shows that glycosylation affects the ECDs of the three NT receptors in different ways too, and that it has roles in shielding and in modulating the receptor conformational ensemble that affect NT binding and may thereby impact NT signalling.

## Supporting Information

Tables S1 and S2, Figures S1-S37, and supporting references.

## Acknowledgments

The authors acknowledge provision of computing resources by the state of Baden-Württemberg through bwHPC and the German Research Foundation (DFG) through grant INST 35/1134-1 FUGG. We thank Stefan Richter for technical support.

## Funding

This research was funded by the European Union’s Horizon 2020 research and innovation programme “Euroneurotrophin” under the Marie Skłodowska-Curie grant agreement No 765704 and the Klaus Tschira Foundation.

## Author contributions

Conceptualization: CA, AT, RCW

Methodology: CA, AT, RCW

Investigation: CA, AT

Visualization: CA, AT

Funding acquisition: RCW

Project administration: RCW

Supervision: RCW

Writing – original draft: CA, AT

Writing – review & editing: CA, AT, RCW

## Competing interests

Authors declare that they have no competing interests. CA is an employee of AstraZeneca at the time of publication and may hold stock options.

## Data availability

Input and coordinate files have been deposited on Zenodo at doi: 10.5281/zenodo.17764984. All other data are available from the authors upon request.

