## Supplementary Information for "Glycosylation-Modulated Conformational Diversity in Neurotrophin Receptors"

### Supporting Information

**Table S1**: Summary of simulated systems showing their glycosylation state and the number and length of the simulation replicas.

| System No | Protein system | Glycosylation state | Replicas | Time/Replica (ns) |
|---|---|---|---|---|
| 1 | p75 dimer – NT3 dimer | + | 3 | 500 |
| 2 | | - | 3 | 500 |
| 3 | p75 monomer – NT3 dimer | + | 3 | 500 |
| 4 | | - | 3 | 500 |
| 5 | p75 monomer | + | 3 | 500 |
| 6 | | - | 3 | 500 |
| 7 | TrkA dimer – NGF dimer | + | 6 | 500 |
| 8 | | - | 6 | 500 |
| 9 | TrkA monomer – NGF dimer | + | 6 | 500 |
| 10 | | - | 6 | 500 |
| 11 | TrkA monomer | + | 3 | 500 |
| 12 | | - | 3 | 500 |
| 13 | TrkB dimer – NT-4/5 dimer | + | 6 | 500 |
| 14 | | - | 6 | 500 |
| 15 | TrkB monomer – NT-4/5 dimer | + | 6 | 500 |
| 16 | | - | 6 | 500 |
| 17 | TrkB monomer | + | 6 | 500 |
| 18 | | - | 6 | 500 |

**Table S2**: N-glycosylation positions on TrkA and TrkB receptors. For TrkB, the exact glycosylation positions are known experimentally.[1,2] For TrkA, the glycosylation positions that are known from the crystal structure (PDB ID: 2IFG[3] ) are shown in normal font, while those that were added based on comparison with TrkB positions are shown in bold font.

| TrkA | TrkB |
|---|---|
| **Asn 67** | Asn 67 |
| Asn 95 | Asn 95 |
| Asn 121 | Asn 121 |
| Asn 188 | Asn 178 |
| **Asn 202** | Asn 205 |
| - | Asn 241 |
| **Asn 253** | Asn 254 |
| Asn 262 | - |
| Asn 281 | Asn 280 |
| - | Asn 338 |
| Asn 358 | - |
| **Asn 401** | Asn 412 |

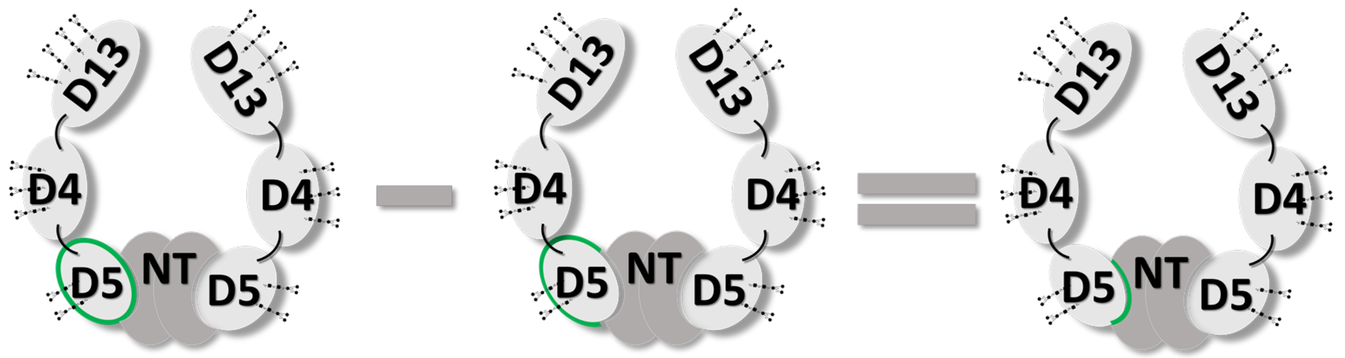


**Figure S1**: Schematic representation of the solvent accessible surface area (SASA) calculations that were performed to estimate the contact area between the extracellular domains of the receptor and the neurotrophin (NT). Here, the contact area between the D5 domain of the Trks and the NT is shown as an example. For this purpose, the SASA of the D5 domain was calculated without (left) and with (middled) the NT present, and the difference between these two values yielded the contact area for D5-NT (right).

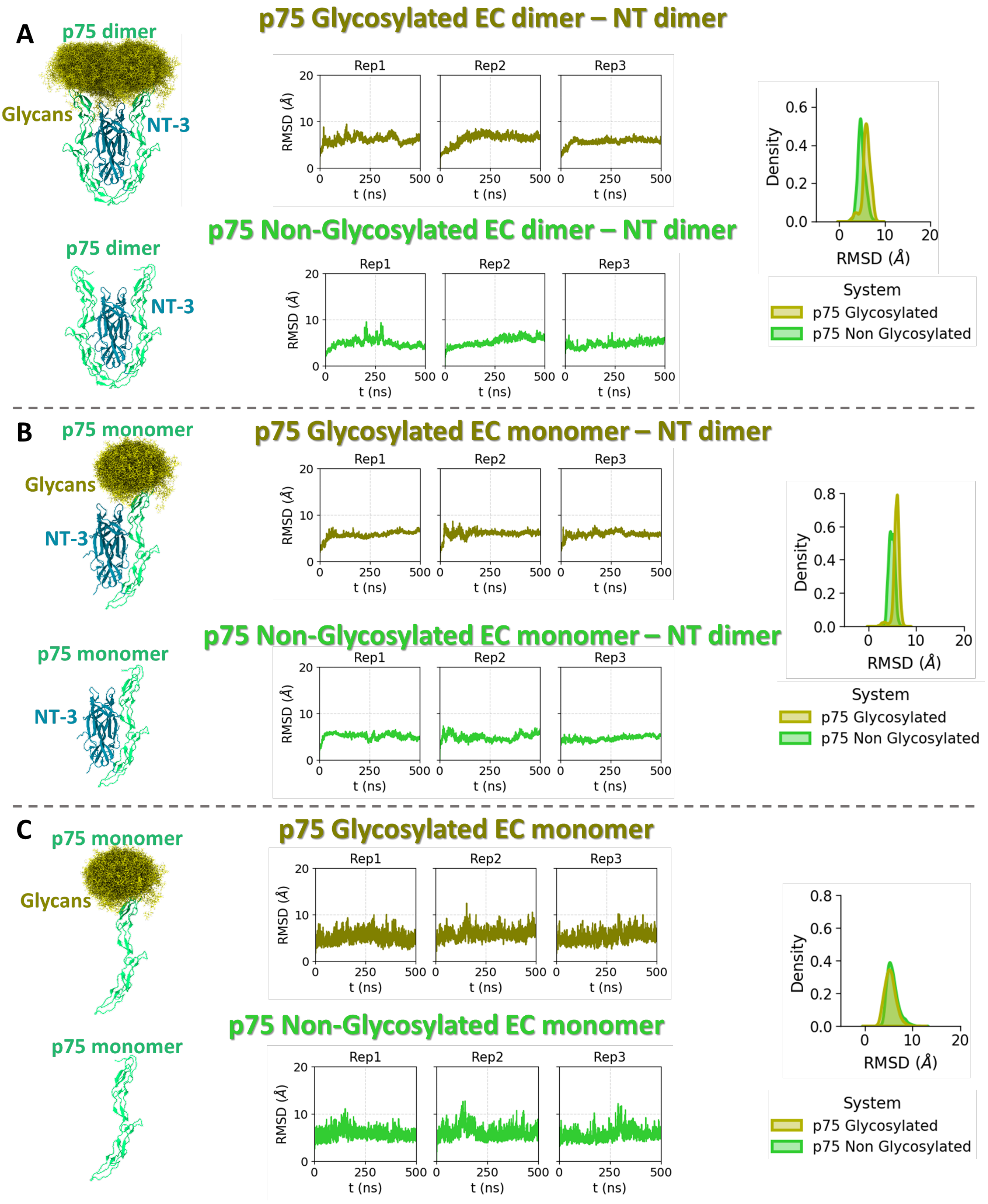


**Figure S2**: RMSD values for the glycosylated and non-glycosylated p75-ECD (A) as a homodimer in complex with NT-3, (B) as a monomer in complex with NT-3 and (C) as a monomer alone. For each system, the time-dependence of the RMSD of the protein backbone compared to the first frame (which corresponds to the crystal structure) is shown for each replica, as well as the distributions of RMSD values across all replicas and all frames of the production runs for the glycosylated and non-glycosylated systems (right-hand plots)

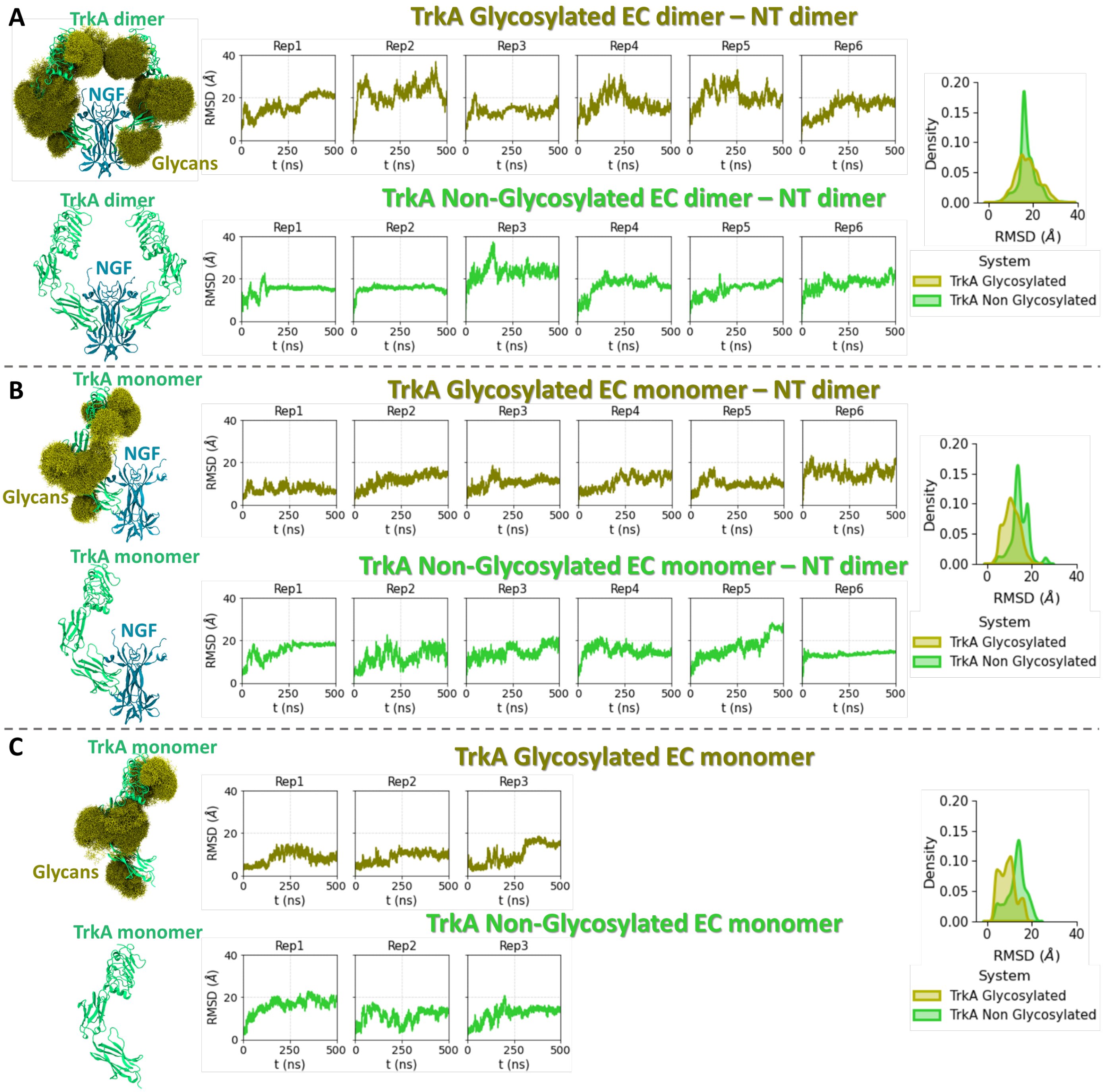


**Figure S3**: RMSD values for the glycosylated and non-glycosylated TrkA systems. (A), (B) and (C) show the RMSD for the TrkA-EC homodimer with the NGF homodimer bound, the TrkA-EC monomer with the NGF homodimer bound, the TrkA-EC monomer systems, respectively, with and without glycans. For each system, the time-dependence of the RMSD of the protein backbone compared to the first frame (which corresponds to the crystal structure) is shown for each replica, as well as the distributions of RMSD values across all replicas and all frames of the production runs for the glycosylated and non-glycosylated systems (right-hand plots).

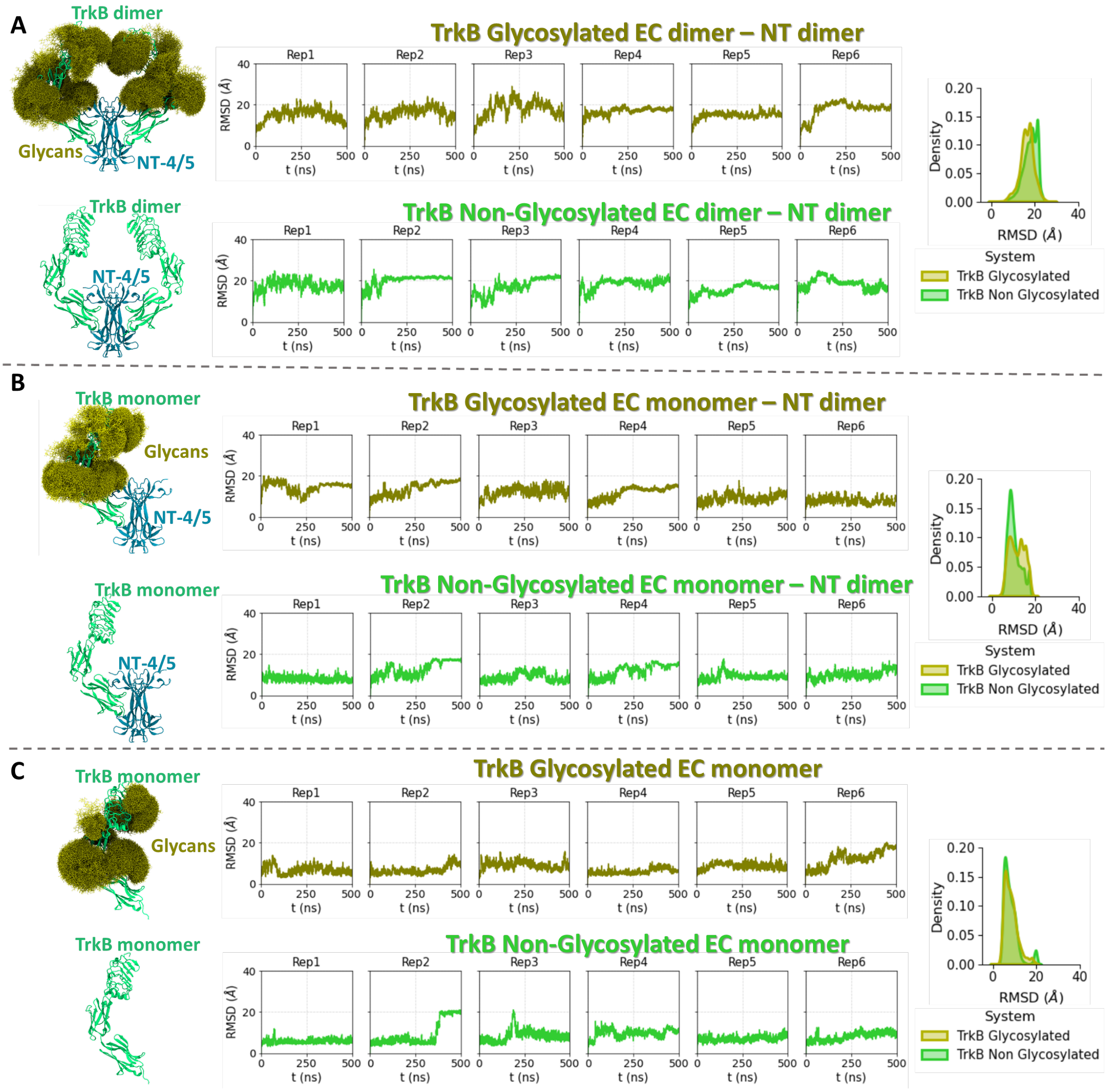


**Figure S4**: RMSD values for the glycosylated and non-glycosylated TrkB systems. (A), (B) and (C) show the RMSD for the TrkB-EC homodimer with the NT-4/5 homodimer bound, the TrkB-EC monomer with the NT-4/5 homodimer bound, the TrkB-EC monomer systems, respectively, with and without glycans. For each system, the time-dependence of the RMSD of the protein backbone compared to the first frame (which corresponds to the crystal structure) is shown for each replica, as well as the distributions of RMSD values across all replicas and all frames of the production runs for the glycosylated and non-glycosylated systems (right-hand plots).

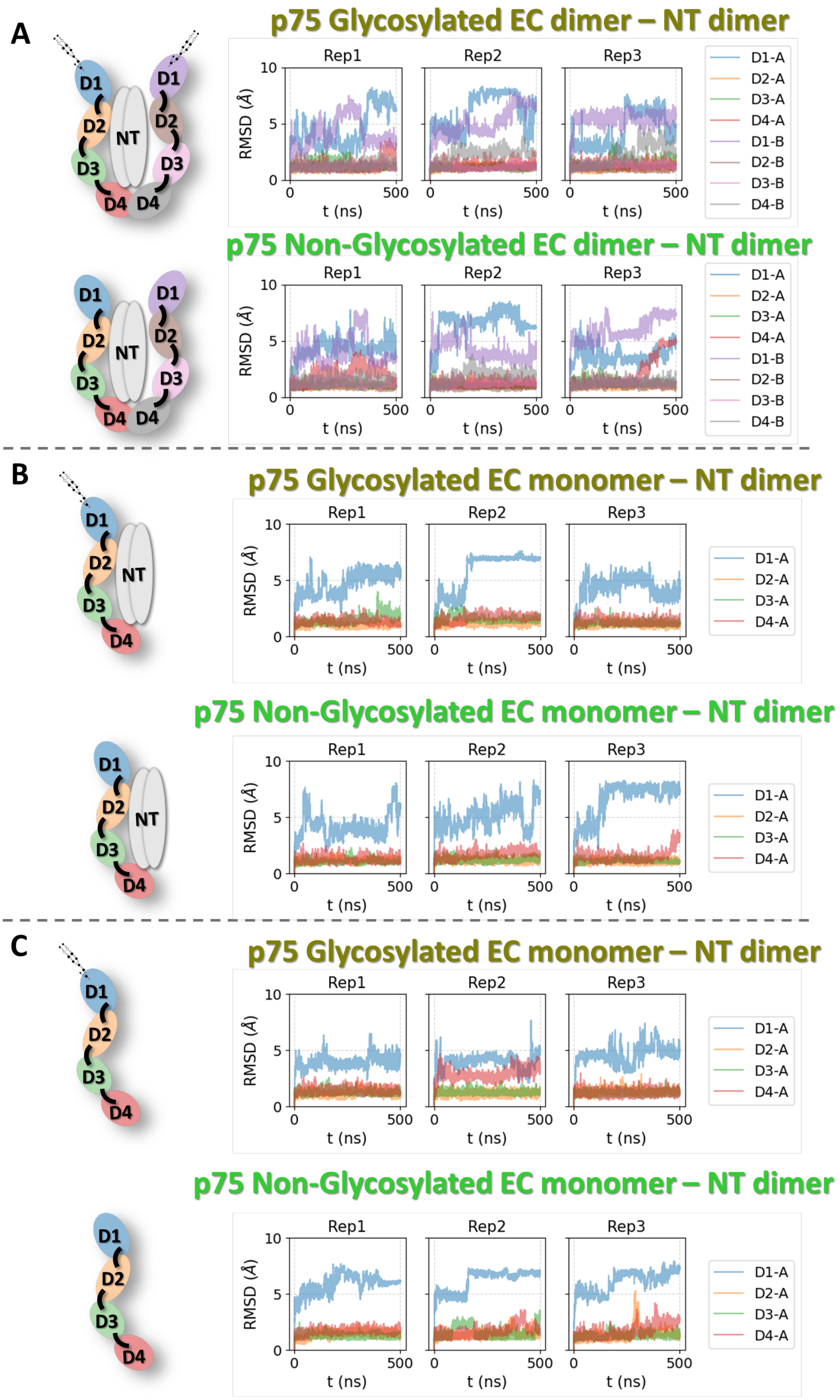


**Figure S5**: RMSD values for each domain for the glycosylated and non-glycosylated p75-ECD (A) as a homodimer in complex with NT-3, (B) as a monomer in complex with NT-3 and (C) as a monomer alone. For each system, the time-dependent RMSD of the domain backbone compared to the first frame is shown for each replica. The RMSD values were calculated after alignment to each domain

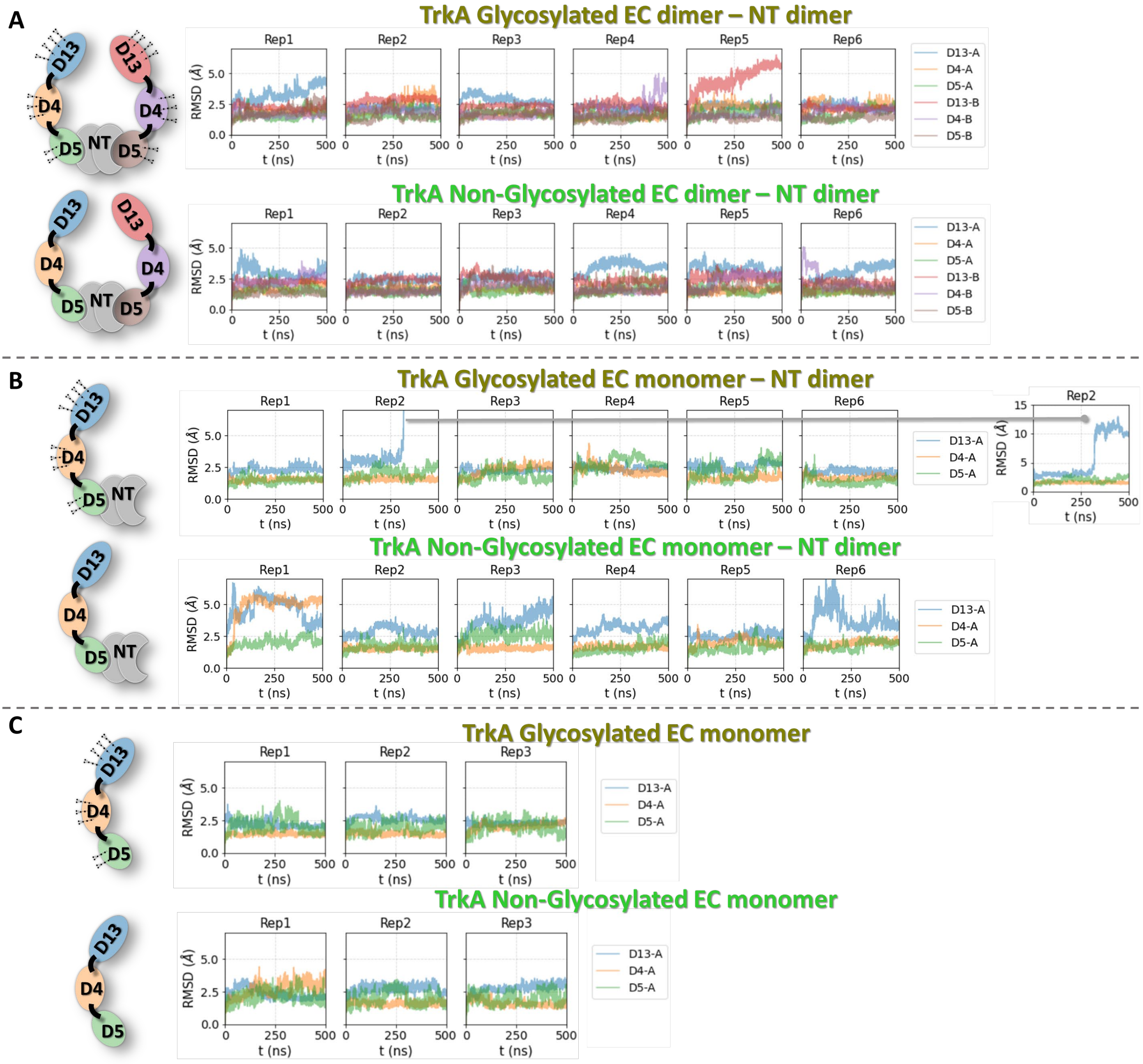


**Figure S6**: RMSD values for each domain of the glycosylated and non-glycosylated TrkA systems. (A), (B) and (C) show the internal RMSD for each domain the TrkA-EC homodimer with the NGF homodimer bound, the TrkA-EC monomer with the NGF homodimer bound, the TrkA-EC monomer systems, respectively, with and without glycans. For each system, the time-dependent RMSD of the domain backbone compared to the first frame is shown for each replica. Before the calculation of the internal RMSD of each domain, alignment of the respective domain at each simulation snapshot was performed. A schematic representation of each system, with color-coding of the domains is shown on the left. The first and second subunits are indicated with A and B letters in the legends.

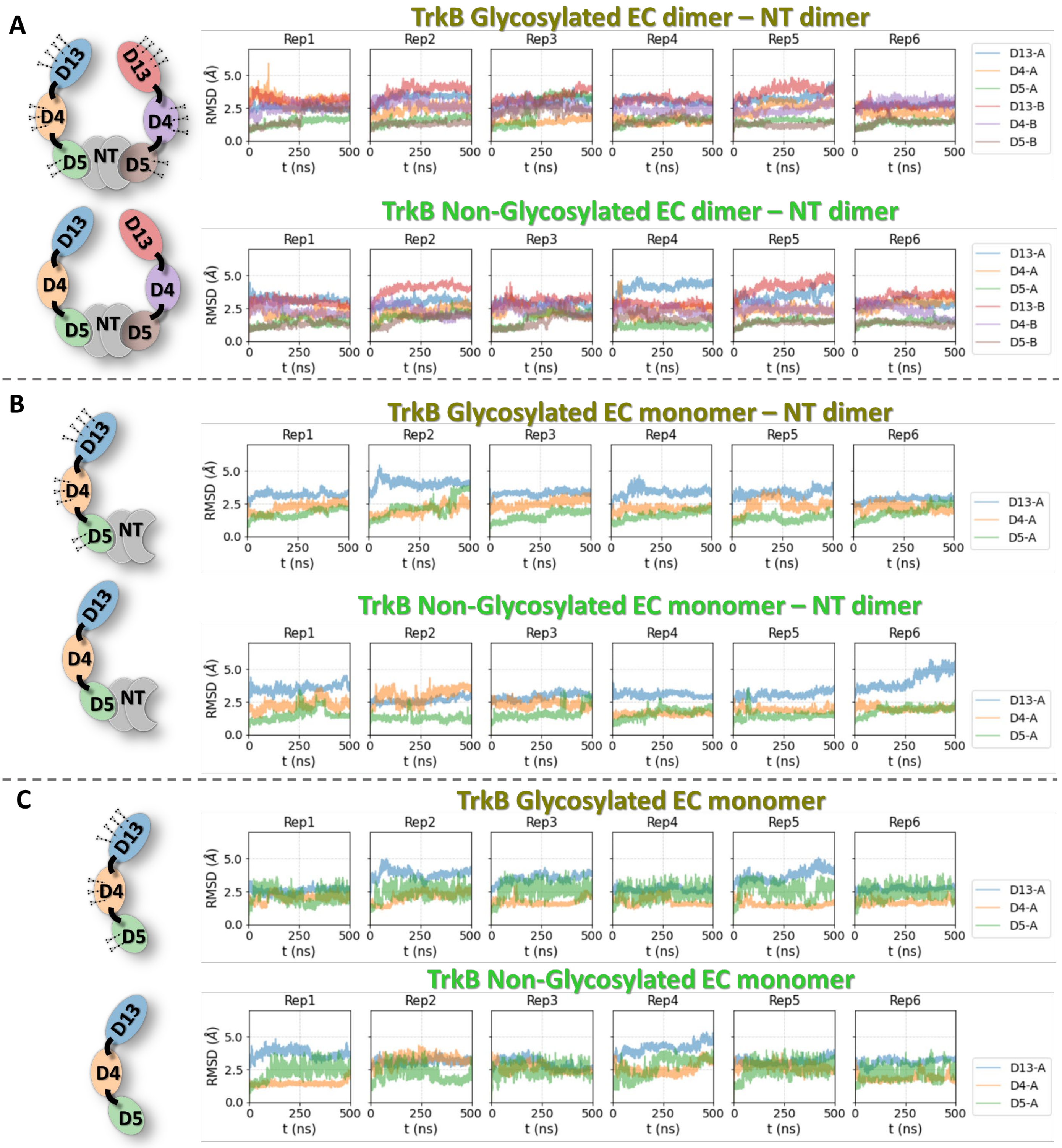


**Figure S7**: RMSD values for each domain of the glycosylated and non-glycosylated TrkB systems. (A), (B) and (C) show the internal RMSD for each domain of the TrkB-EC homodimer with the NT-4/5 homodimer bound, the TrkB-EC monomer with the NT-4/5 homodimer bound, the TrkB-EC monomer systems, respectively, with and without glycans. For each system, the time-dependent RMSD of the domain backbone compared to the first frame is shown for each replica. Before the calculation of the internal RMSD of each domain, alignment of the respective domain at each simulation snapshot was performed. A schematic representation of each system, with color-coding of the domains is shown on the left. The first and second monomers are indicated by A and B letters in the legends.

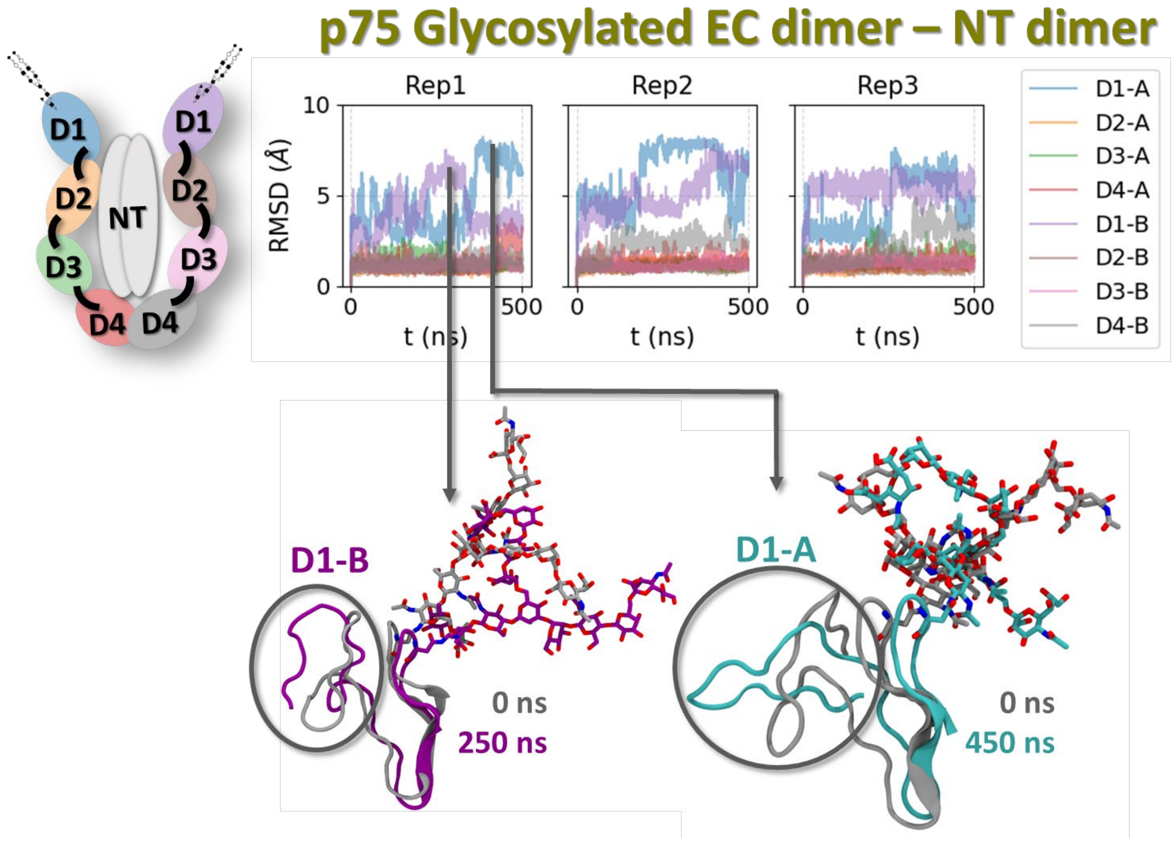


**Figure S8**: Closer examination of the p75-D1 domain conformations with the glycan at Asn 60, which is covalently attached to the D1 domain. A schematic representation of the simulated system, with color-coding of the domains is shown on the left. Domain-based RMSD vs time plots for three replica simulations show the highest RMSD values for D1 domains. Two example structures with high RMSD values are shown with the D1-glycan aligned to the input structure from the crystal structure. The high RMSD conformations are shown in purple and cyan for the two D1 monomers and the initial structures are shown in grey. The D1 domain is shown in cartoon representation, while the glycan is shown in stick representation.

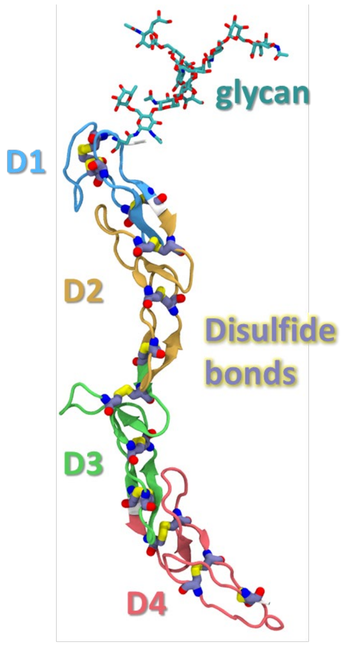


**Figure S9**: Disulfide bond ladder in the structure of the p75-ECD (PDB ID: 3BUK). The D1-D4 domains are shown in cartoon representation. The disulfide bonds are shown in licorice representation.

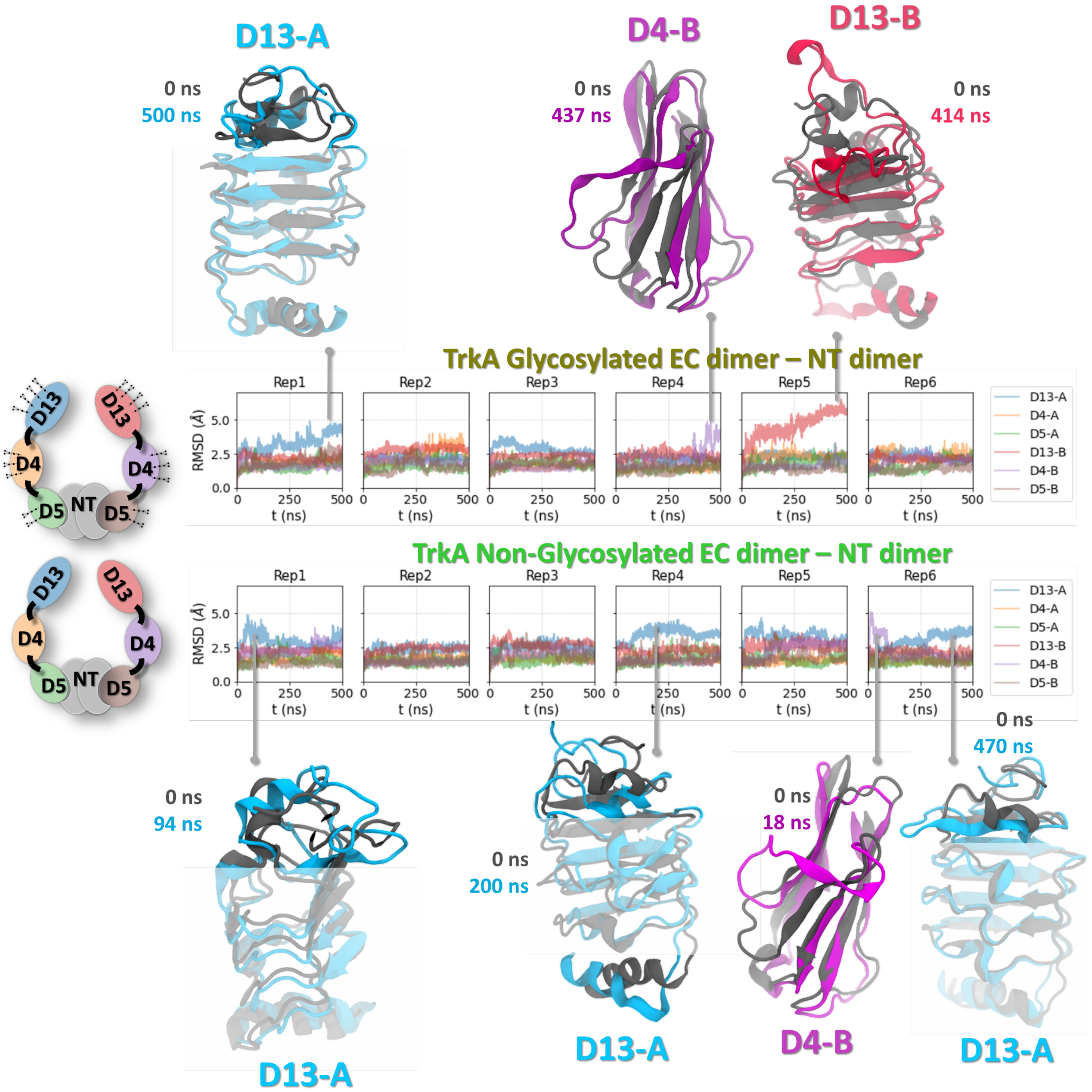


**Figure S10**: RMSD values for each domain of the glycosylated and non-glycosylated TrkA-ECD homodimer bound to NGF. For each system, the time-dependent RMSD of the protein domain backbone compared to the first frame is shown for each replica. Before the calculation of the internal RMSD of each domain, an alignment of the respective domain at each simulation snapshot was performed. A schematic representation of each system, with color-coding of the domains is shown on the left. For cases with high RMSD values, the conformations of the domains are shown in cartoon representation aligned to the first frame.

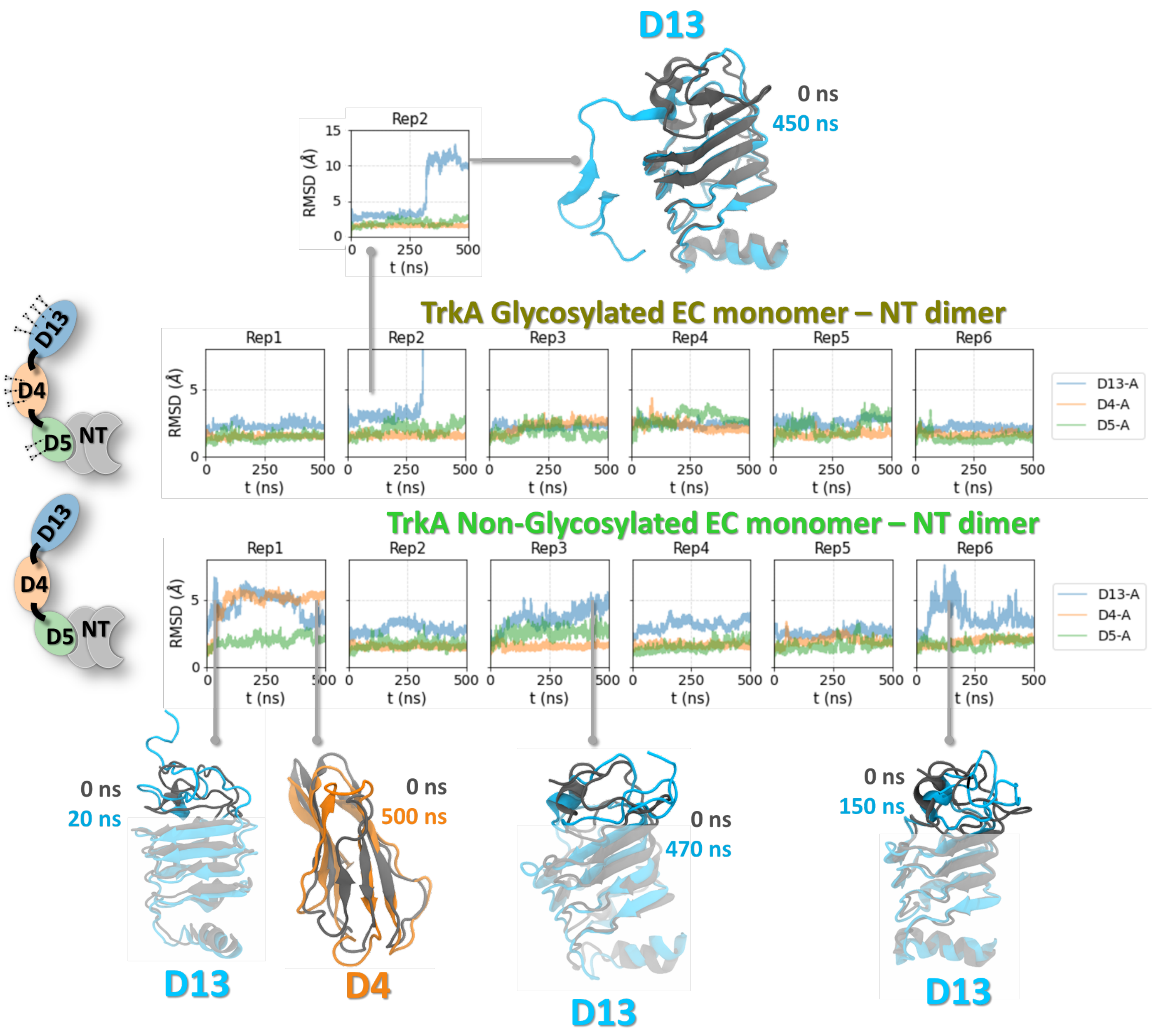


**Figure S11**: RMSD values for each domain glycosylated and non-glycosylated TrkA-EC monomer bound to NGF system. For each system, the time-dependent RMSD of the protein domain backbone compared to the first frame is shown for each replica. Before the calculation of the internal RMSD of each domain, alignment of the respective domain at each simulation snapshot was performed. A schematic representation of each system, with color-coding of the domains is shown on the left. For cases with high RMSD values, the conformations of the domains are shown in cartoon representation aligned to the first frame.

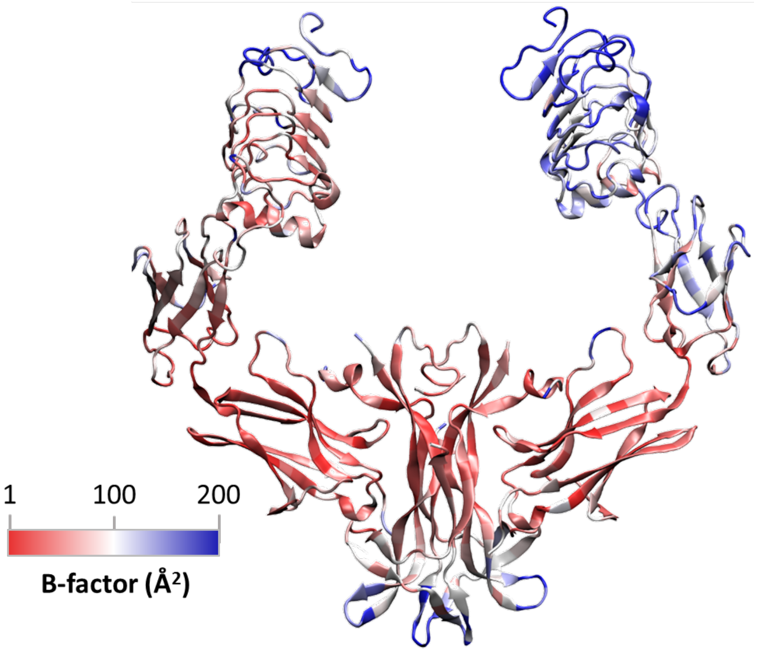


**Figure S12**: B-factors of the TrkA-EC bound to NGF from the crystal structure (PDB ID: 2IFG).[3]

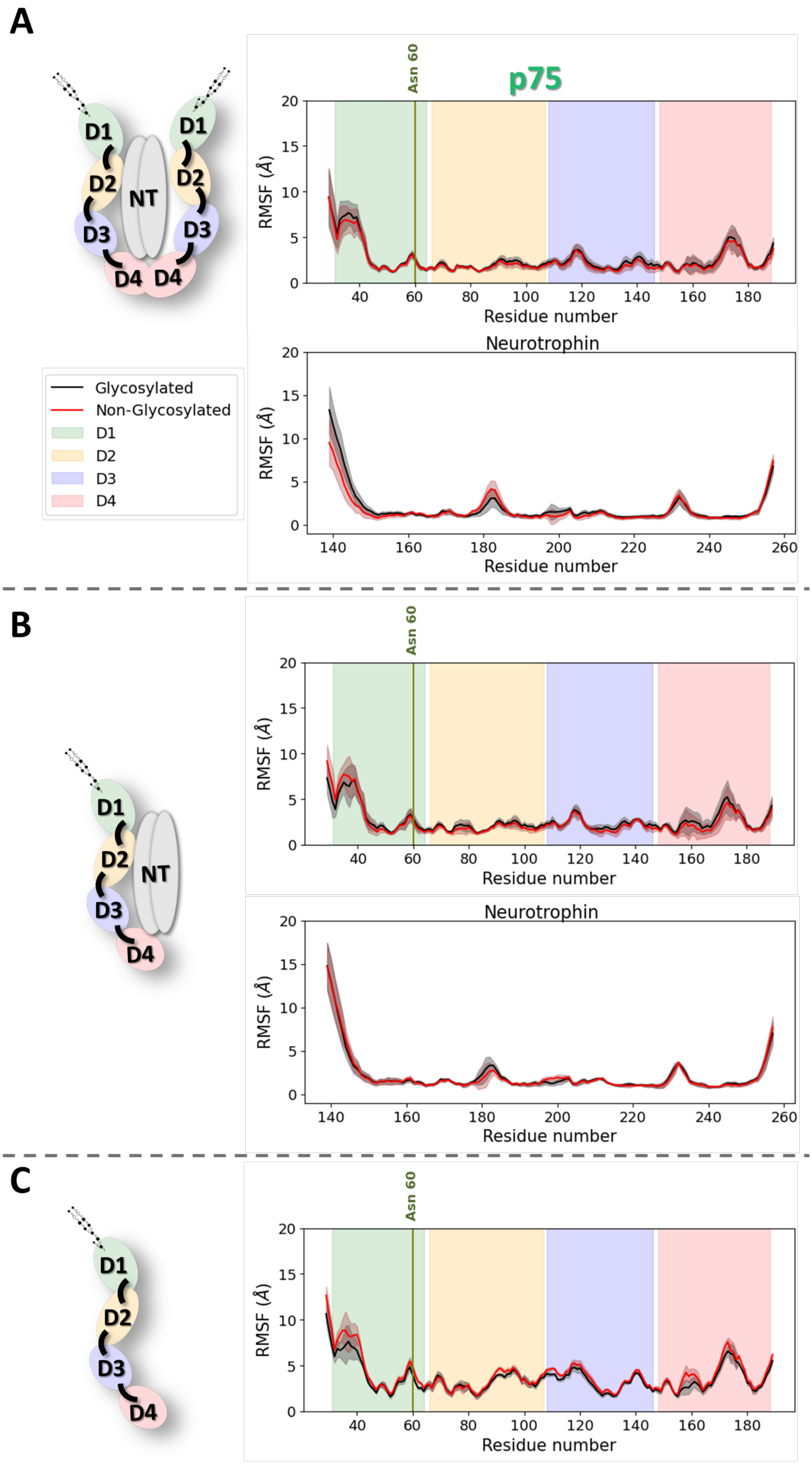


**Figure S13**: RMSF values for the p75-ECD (A) as a homodimer in complex with NT-3, (B) as a monomer in complex with NT-3, and (C) as a monomer alone, with (black) and without (red) glycosylation. The RMSF for each residue was calculated as an average from the three replica simulations and the standard deviation at each point is shown with shadowed areas. The different domains of p75-EC are color-coded and the glycosylated Asn 60 is indicated with a vertical line.

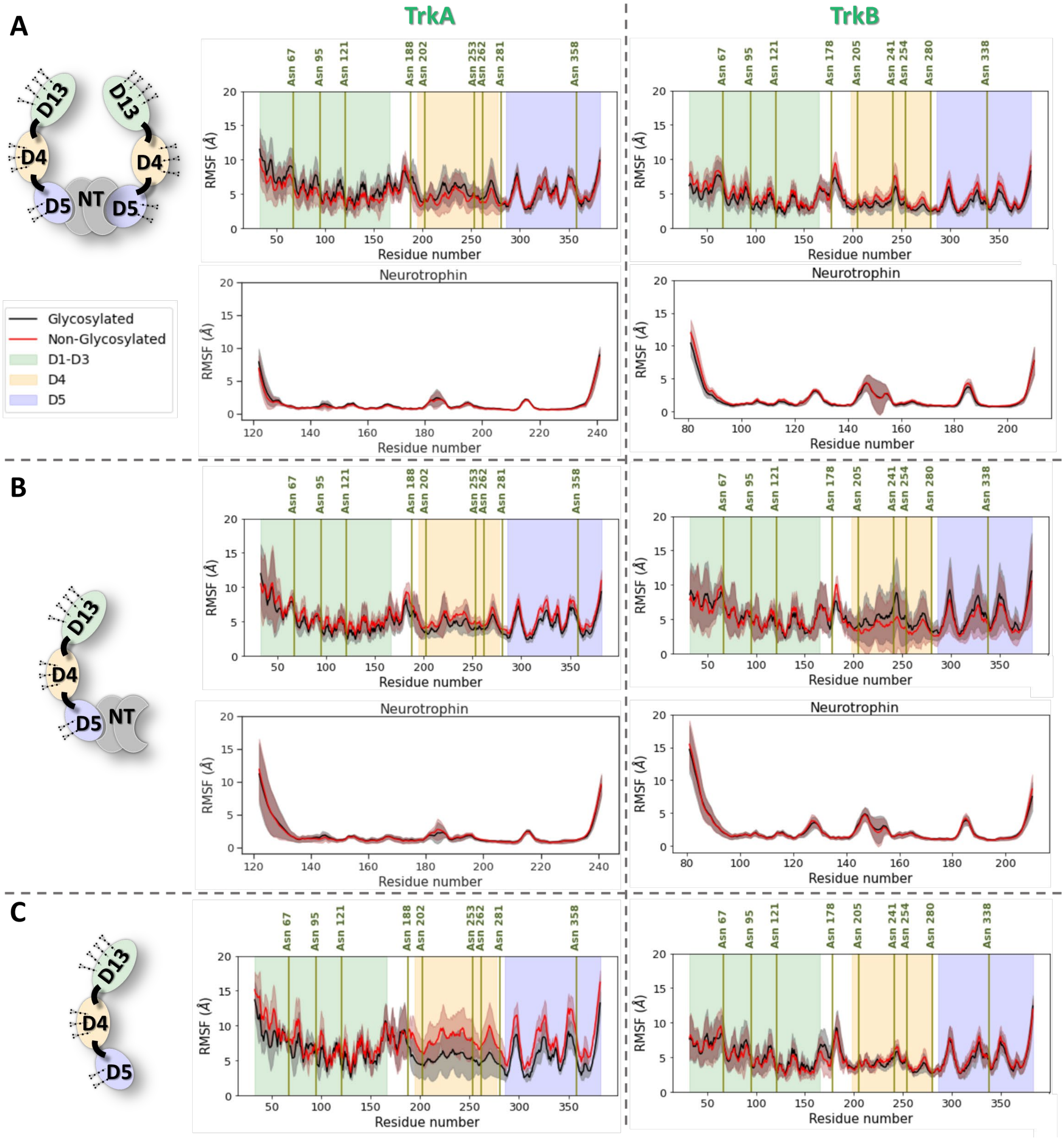


**Figure S14**: RMSF values for all the glycosylated and non-glycosylated TrkA and TrkB systems. (A), (B) and (C) show the average RMSF values from all replica simulations for each receptor and NT residue of the TrkA/B-EC homodimer with the NGF/NT-4/5 homodimer bound, the TrkA/B-EC monomer with the NGF/ NT-4/5 homodimer bound, and the TrkA/B-EC monomer systems, respectively, with or without glycans. The average RMSF values from all replica simulations are shown for each residue in black and red lines for the glycosylated and non-glycosylated systems, respectively. The standard deviation for each residue is shown with shadowed areas. The positions of glycosylated asparagines are shown with vertical lines. Before the RMSF calculation, the systems were aligned to an average structure from the simulations.

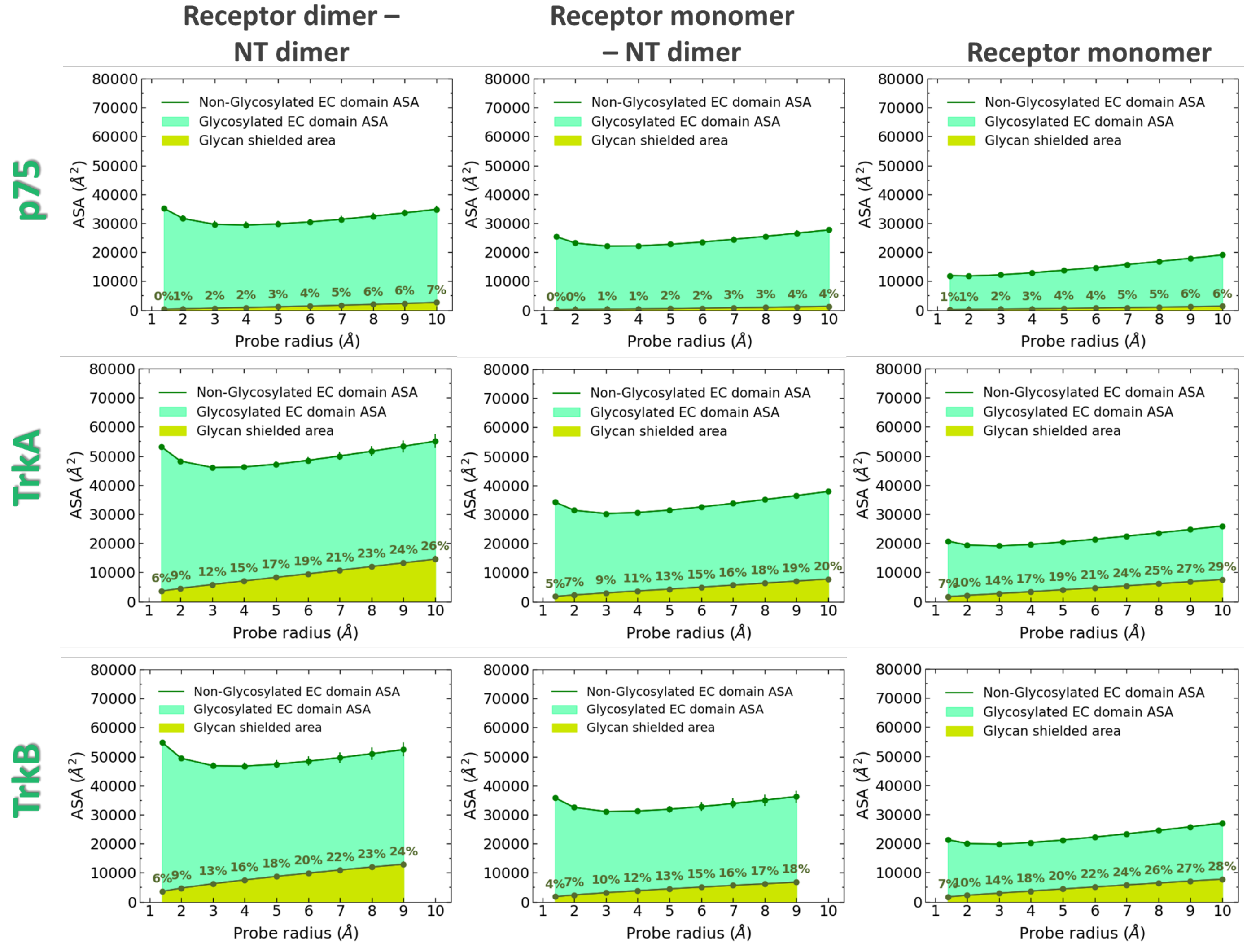


**Figure S15**: Accessible surface area (ASA) of the p75, TrkA and TrkB systems and the area shielded by glycans at multiple probe radii from 1.4 Å (water molecule) to 10 Å (protein-sized molecule). The values were averaged across all replicas. The area shielded by the glycans is presented in yellow (rounded % values are reported), whereas the green line represents the accessible area of the protein in the absence of glycans. Highlighted in green is the area that remains accessible in the presence of glycans.

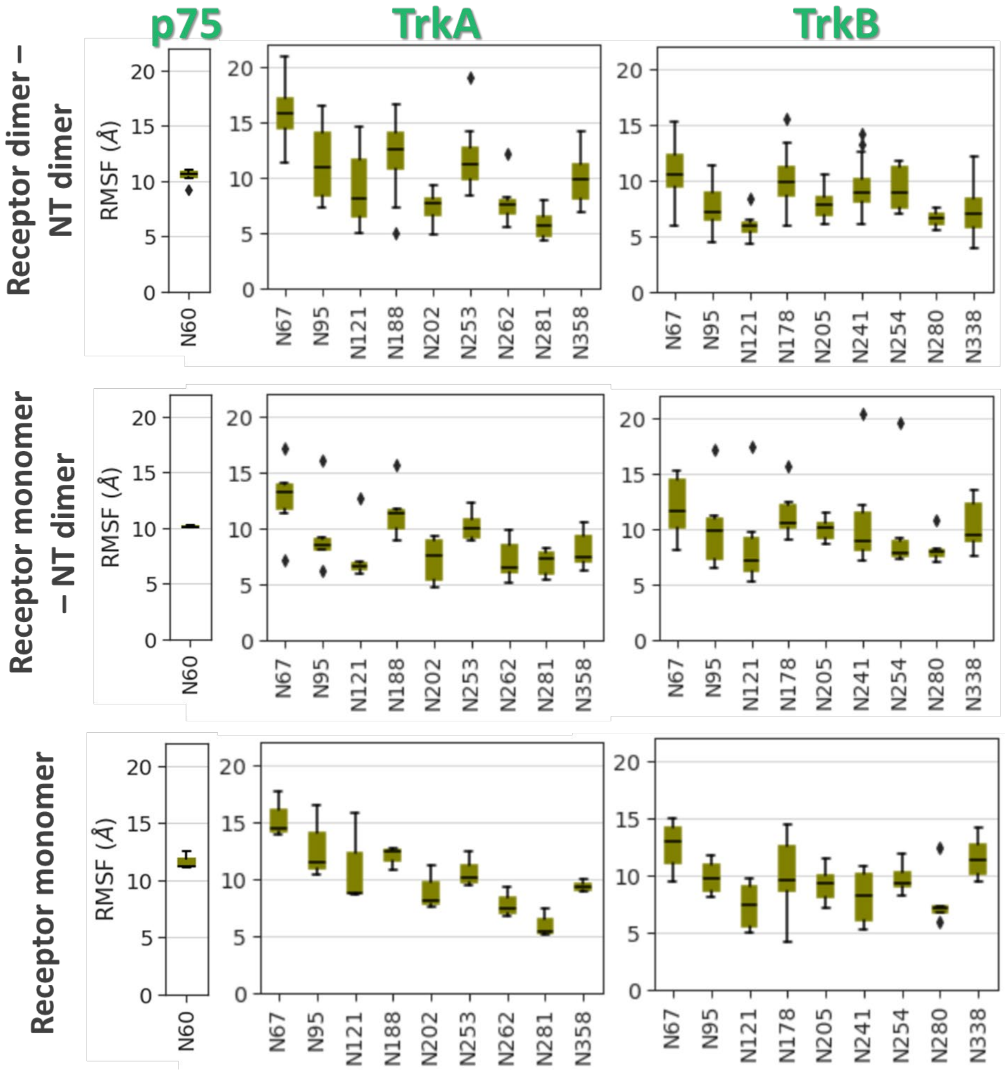


**Figure S16**: Distributions of the RMSF values of all glycans attached to asparagine residues in all the p75, TrkA and TrkB glycosylated systems. Before the RMSF calculation, alignment of each EC segment to its initial structure (frame 0) was performed.

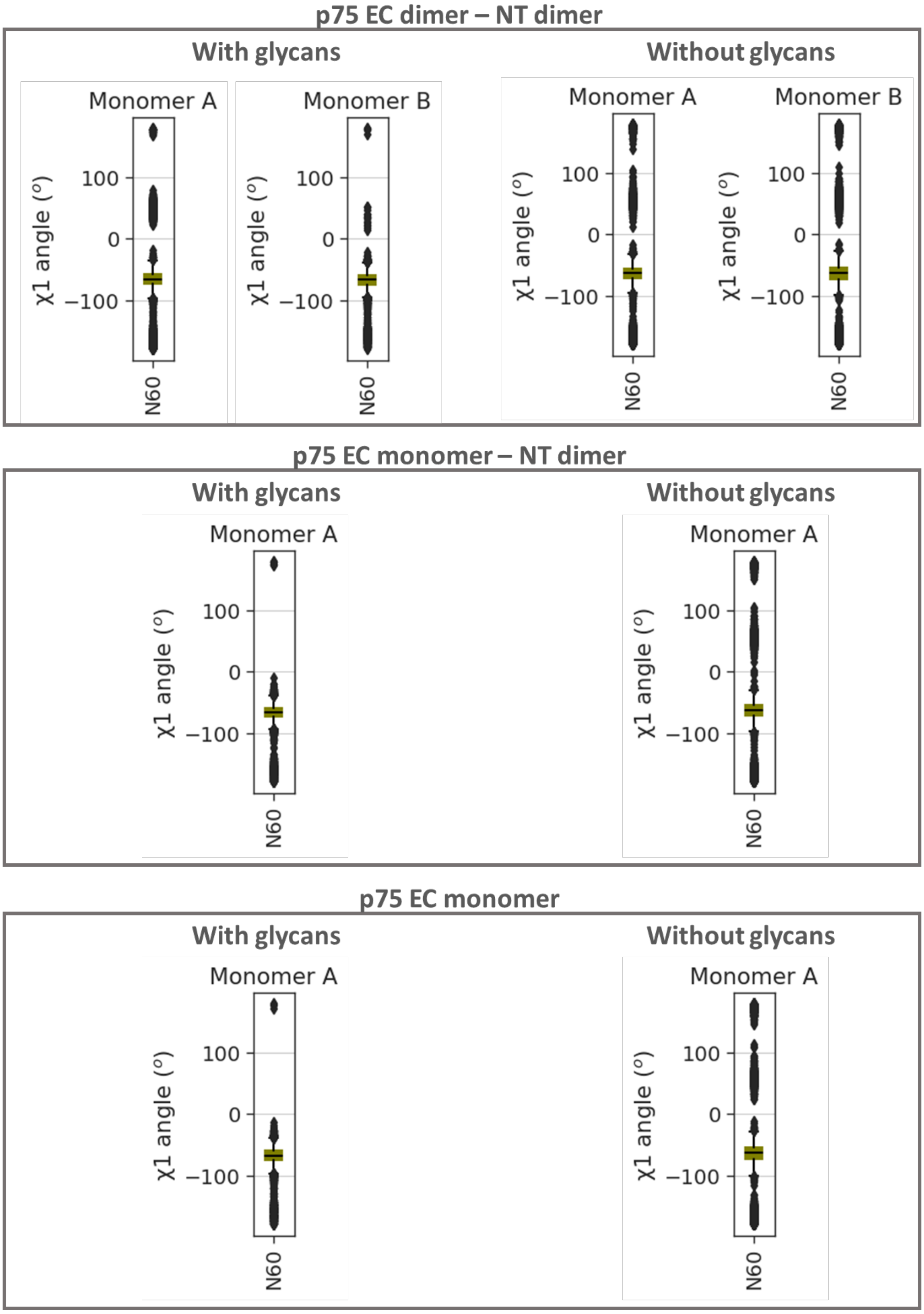


**Figure S17**: Distributions of the χ1 dihedral angle of N60 for the three p75-ECD systems with and without glycosylation.

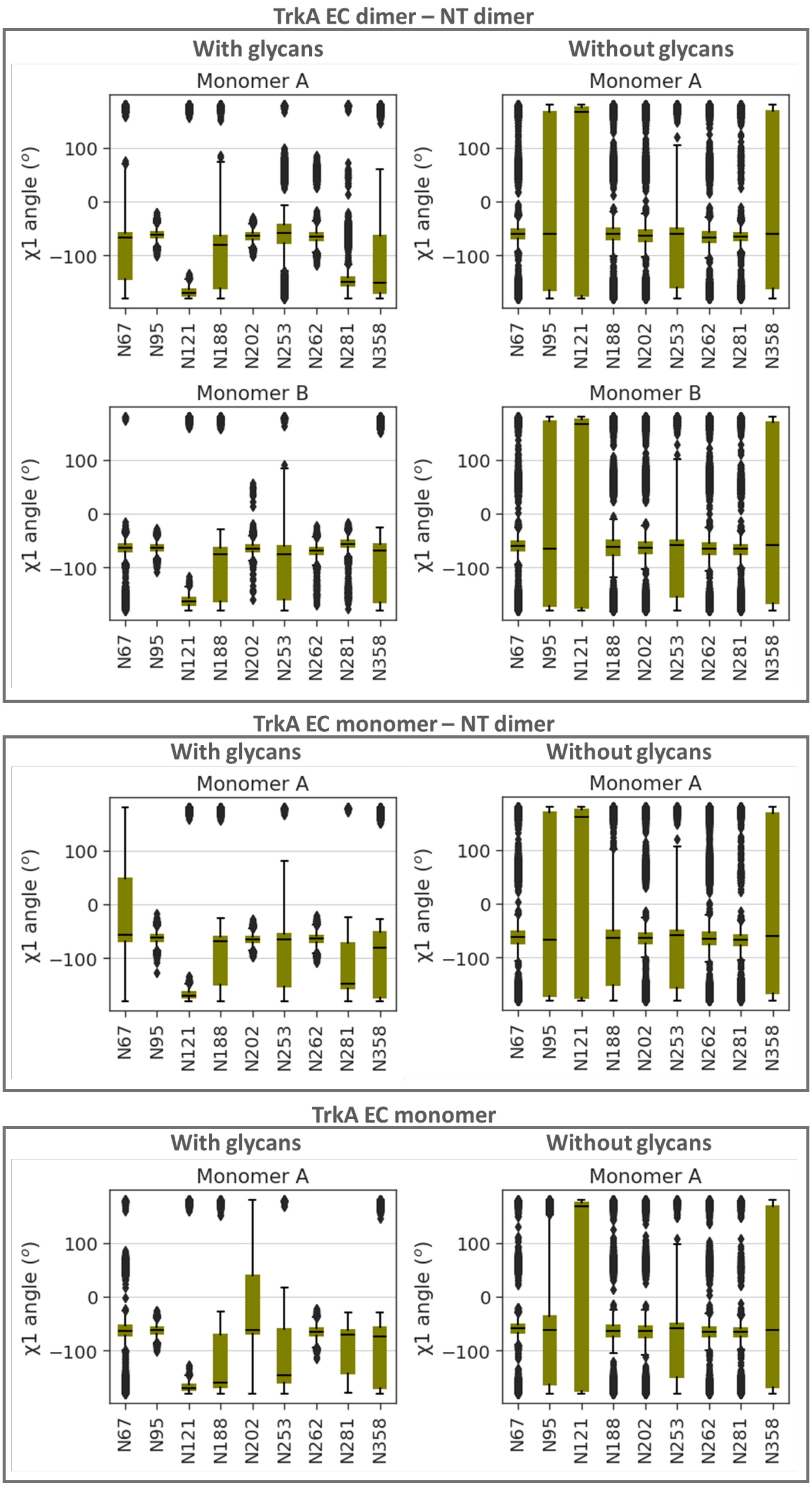


**Figure S18**: Distributions of the χ1 dihedral angle of glycosylated Asn residues in TrkA. A comparison of the χ1 distributions in the glycosylated and non-glycosylated TrkA systems is shown.

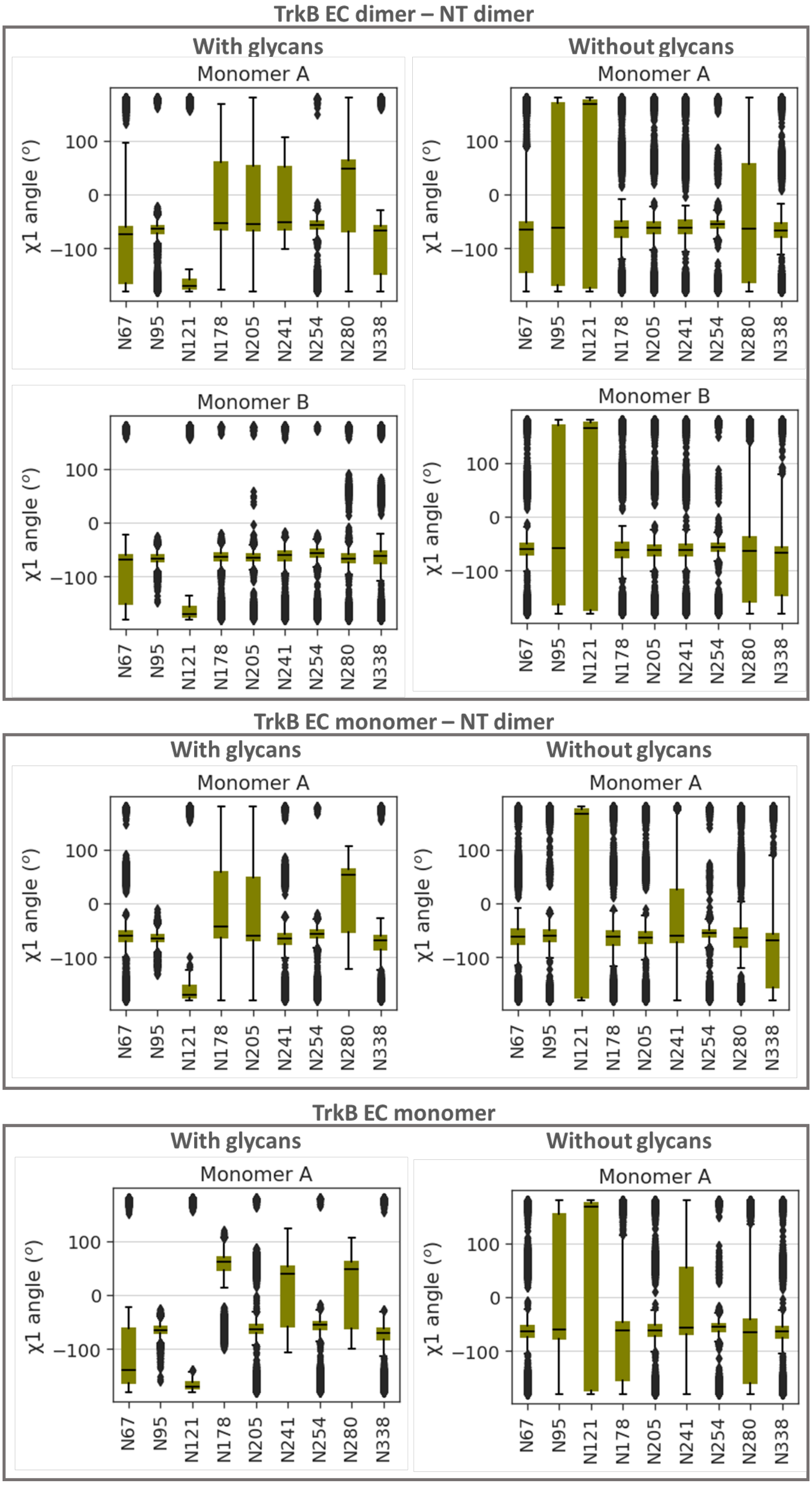


**Figure S19**: Distributions of the χ1 dihedral angle of glycosylated Asn residues in TrkB. A comparison of the χ1 distributions in the glycosylated and non-glycosylated TrkB systems is shown.

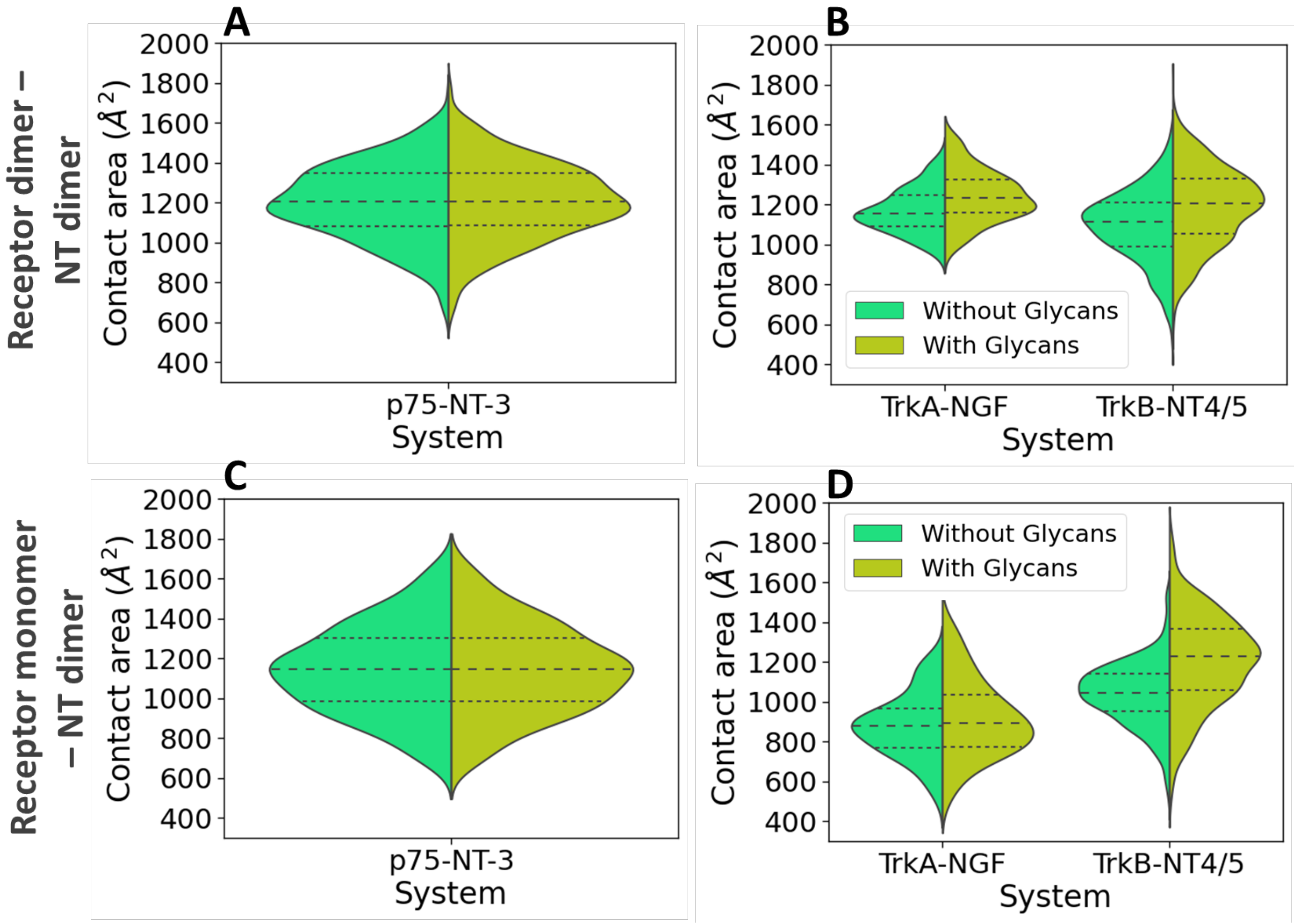


**Figure S20**: Distributions of the contact area between the extracellular domains of the p75, TrkA and TrkB receptors with the NTs. The systems with the receptor dimer ECD bound to NTs with and without glycans are shown in (A) and (B), while the systems with the receptor monomer ECDs bound to NTs with and without glycans are shown in (C) and (D). For the systems with the ECD dimers, the contact area for each monomer was calculated separately.

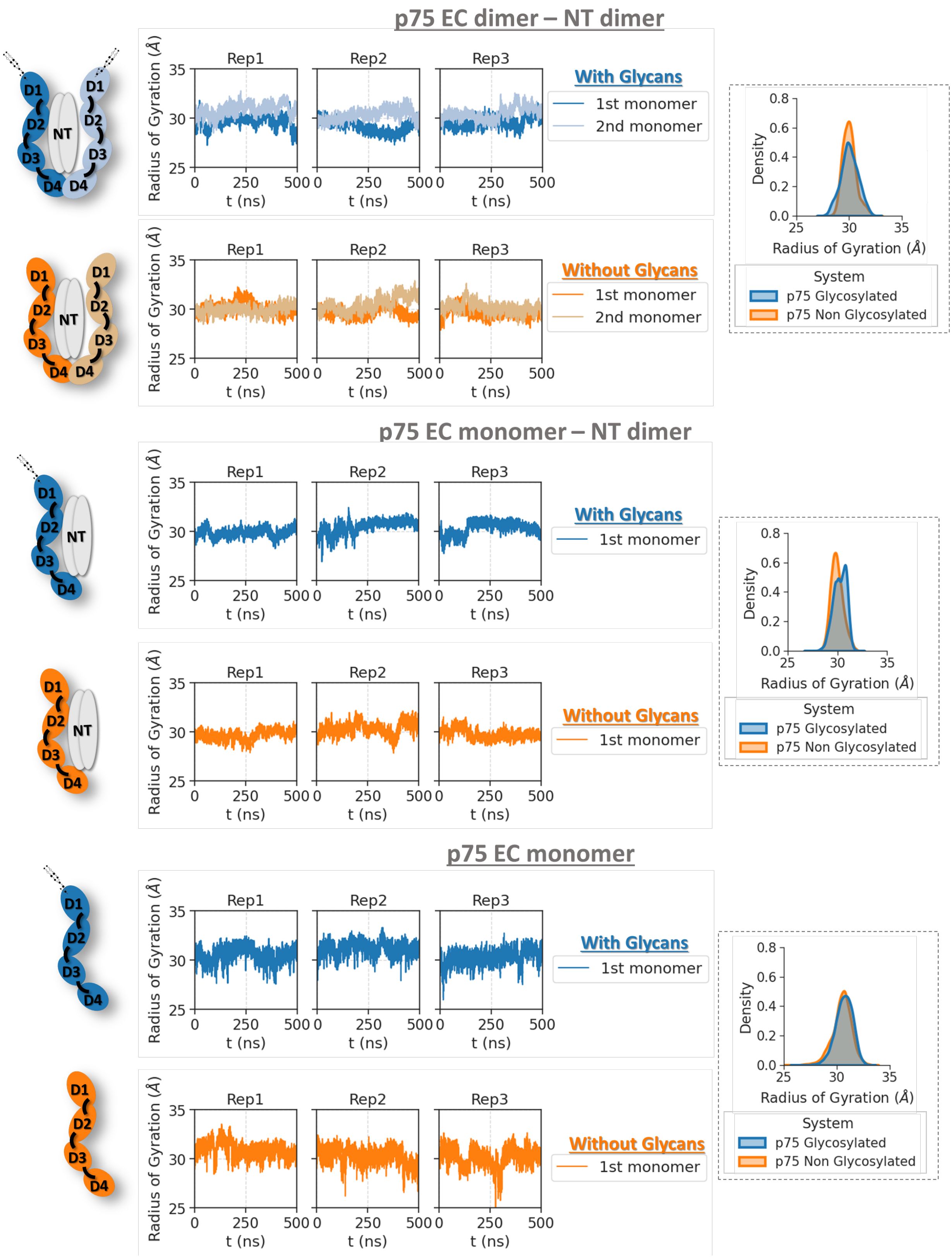


**Figure S21**: Time evolution and distribution of the radius of gyration for the systems of the p75-ECD as a homodimer in complex to NT-3, as a monomer in complex with NT-3, and as a monomer alone, with (blue) and without (orange) glycans. The color-coding of each domain is shown in the schematic representations of the systems on the left of the plots. For the system of the p75-ECD as a homodimer in complex with NT-3, the radius of gyration was calculated for each p75 domain separately.

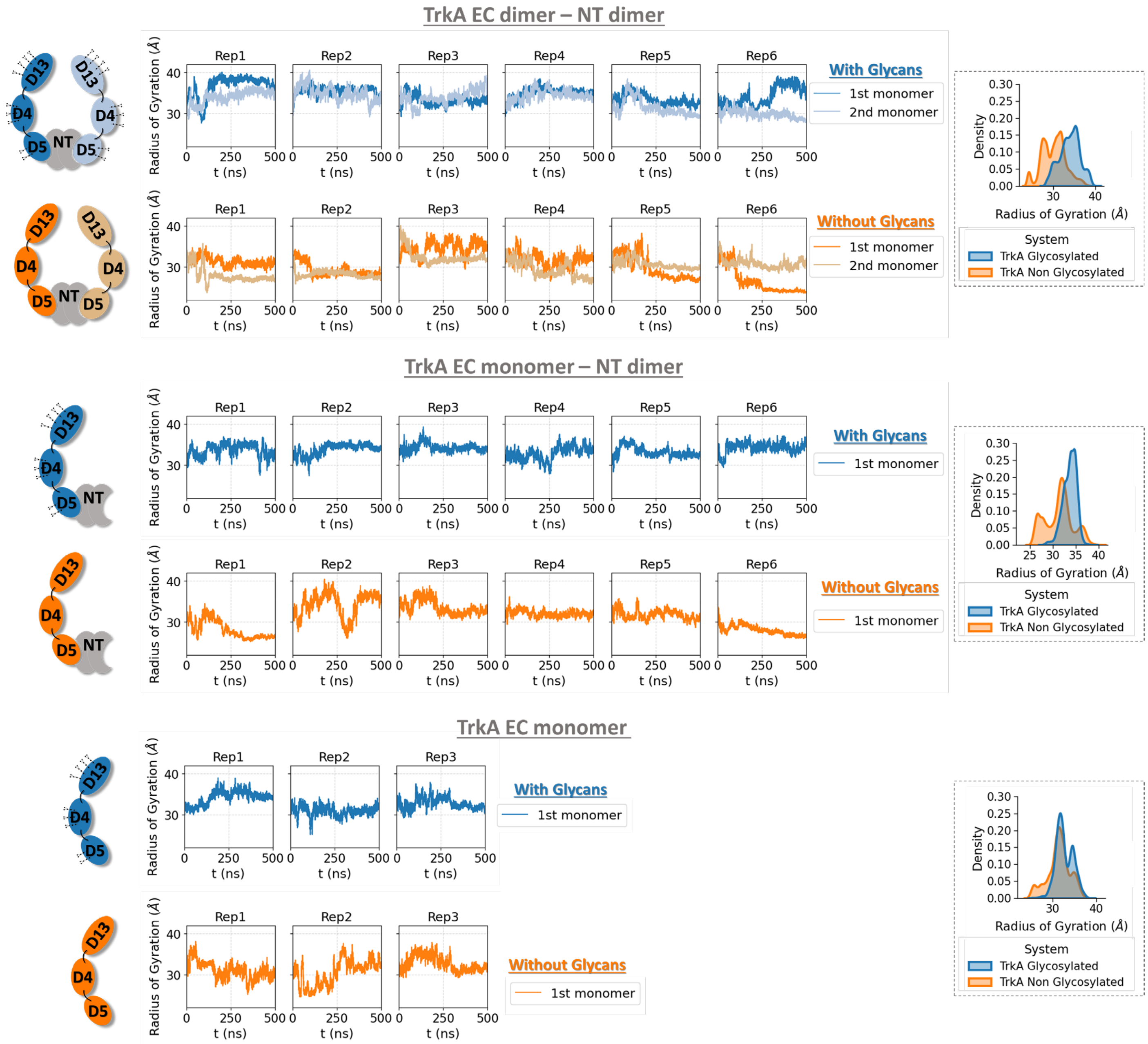


**Figure S22**: Time evolution and distribution of the radius of gyration of the receptor backbone of the glycosylated and non-glycosylated TrkA systems. The color-coding of each domain is shown in the schematic representations of the systems on the left of the plots. For the homodimeric system, the radius of gyration was calculated for each Trk domain separately.

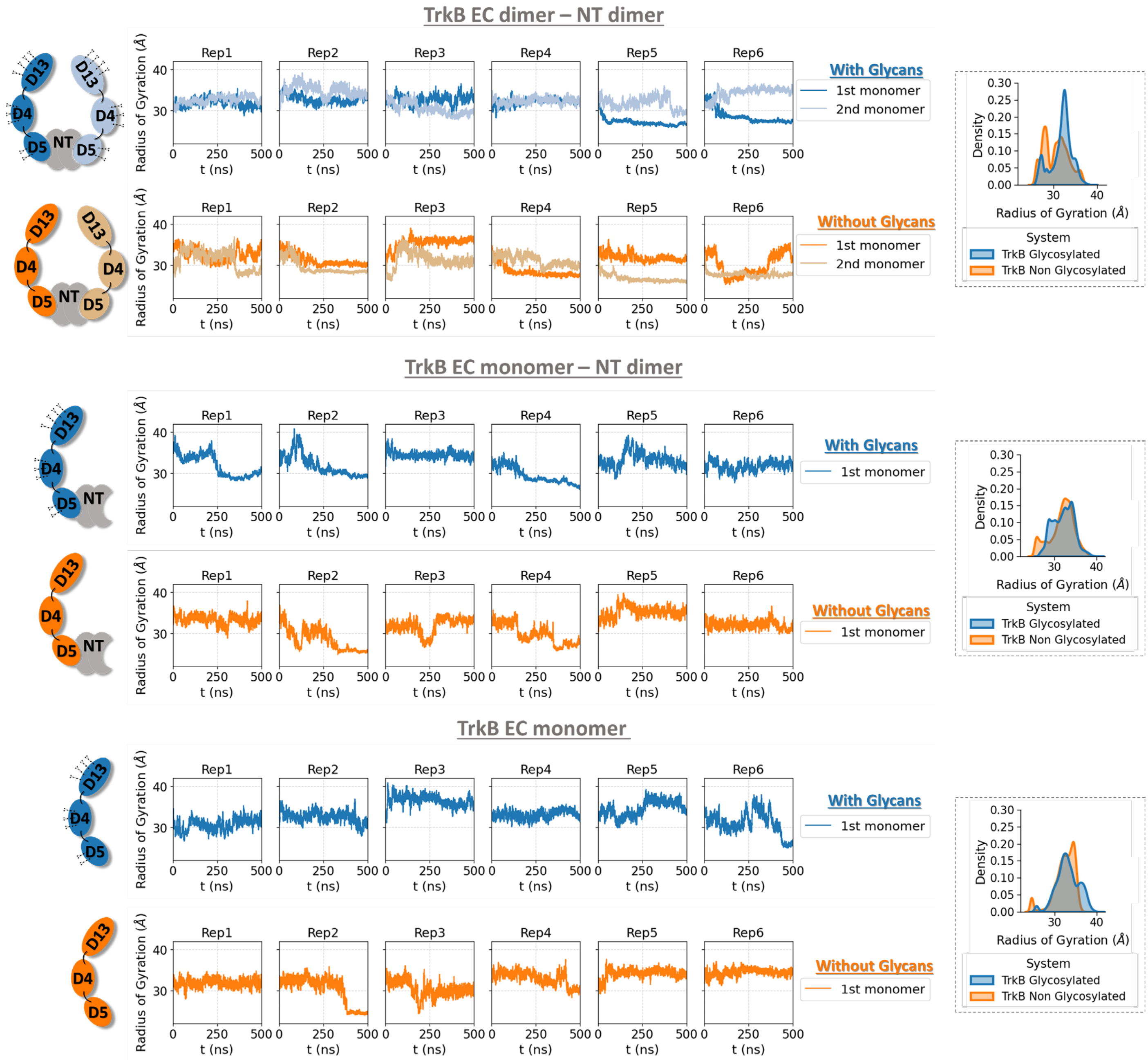


**Figure S23**: Time evolution and distribution of the radius of gyration of the receptor backbone of the glycosylated and non-glycosylated TrkB systems. The color-coding of each domain is shown in the schematic representations of the systems on the left of the plots. For the homodimeric system, the radius of gyration was calculated for each Trk domain separately.

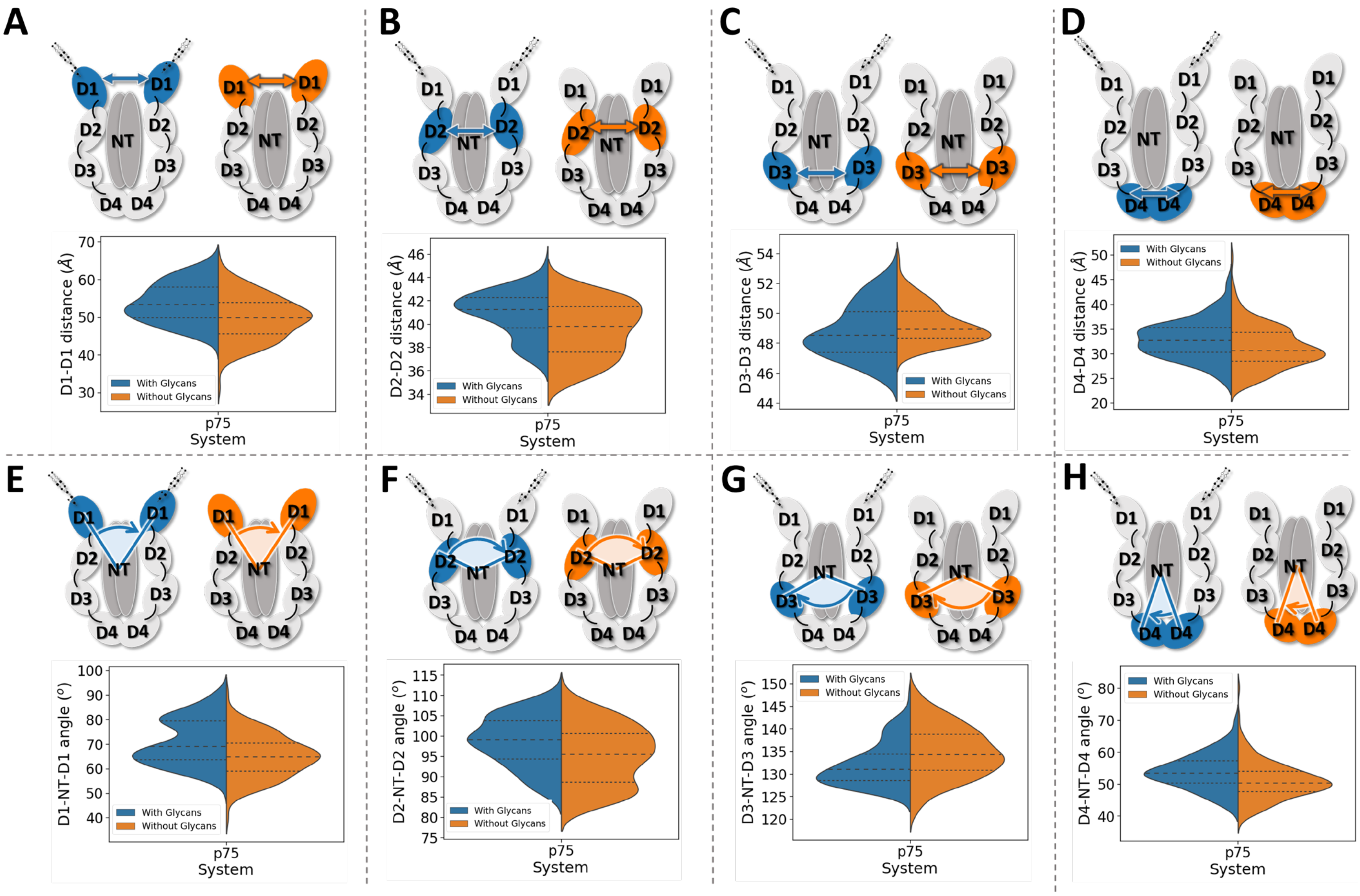


**Figure S24:** Distributions of the distances and angles between the four p75 extracellular domains. (A)-(C) Distances between the COGs of the D1-D1, D2-D2, D3-D3 and D4-D4 domains, in the presence (blue) and absence (orange) of glycosylation. (E)-(H) Angles defined by the COGs of the D1-NT-D1, D2-NT-D2, D3-NT-D3 and D4-NT-D4 domains. The distances and angles were calculated every 100 frames and the final distributions from the glycosylated and non-glycosylated p75 systems are shown in violin plots.

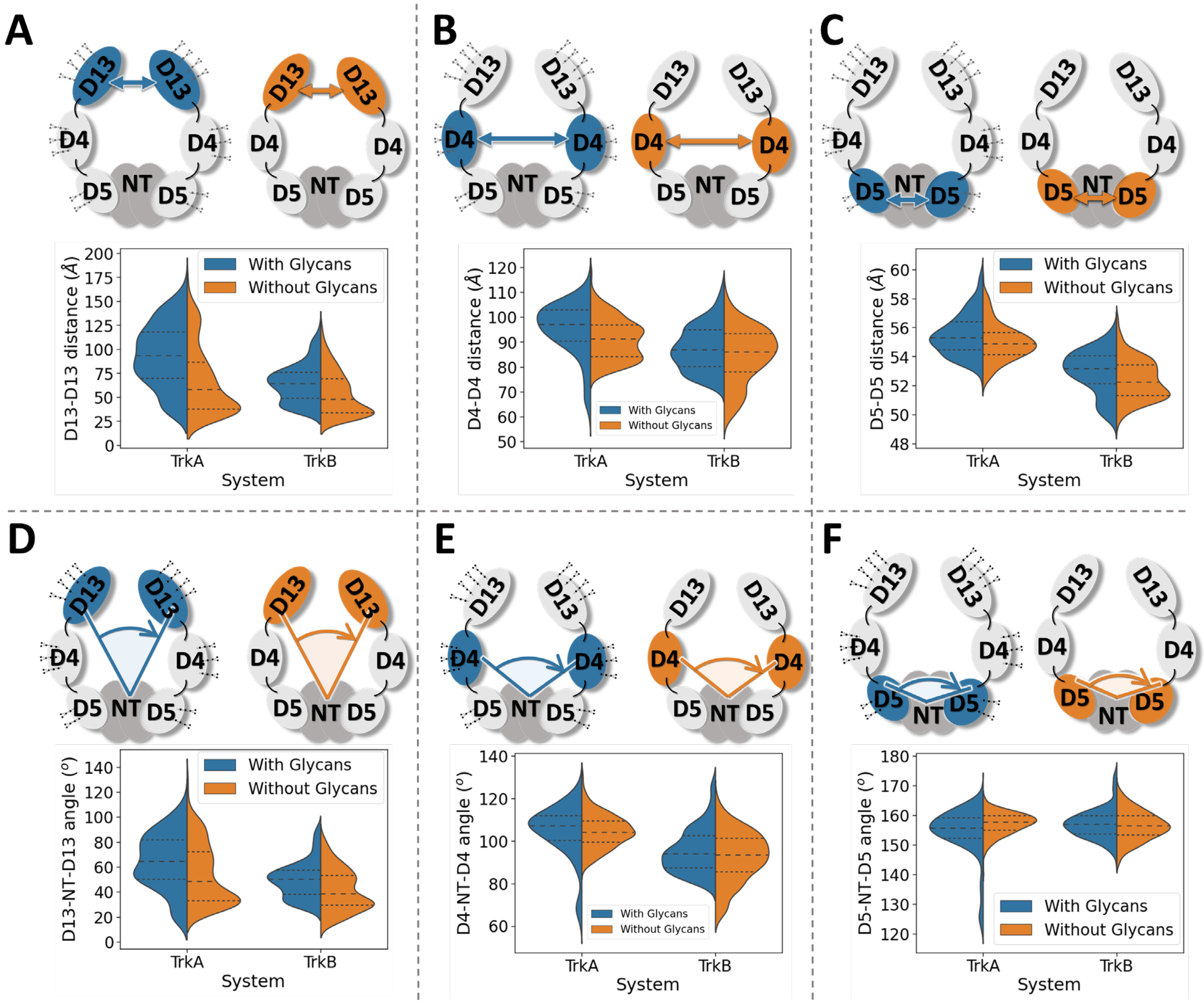


**Figure S25:** Distributions of the distances and angles between the Trk receptor monomer domains. (A, B, C) Calculated distances between the centers of geometry (COG) of the D13, D4 and D5 domains of the two different receptor monomers. (D, E, F) Calculated angles between the COG of one of the same domains from the first receptor monomer, the COG of the NT and the COG of the same domain from the second monomer. The distances and angles were calculated every 100 frames and the final distributions from the glycosylated and non-glycosylated TrkA and TrkB systems are shown in violin plots. The quartiles of the distributions are indicated with dashed lines.

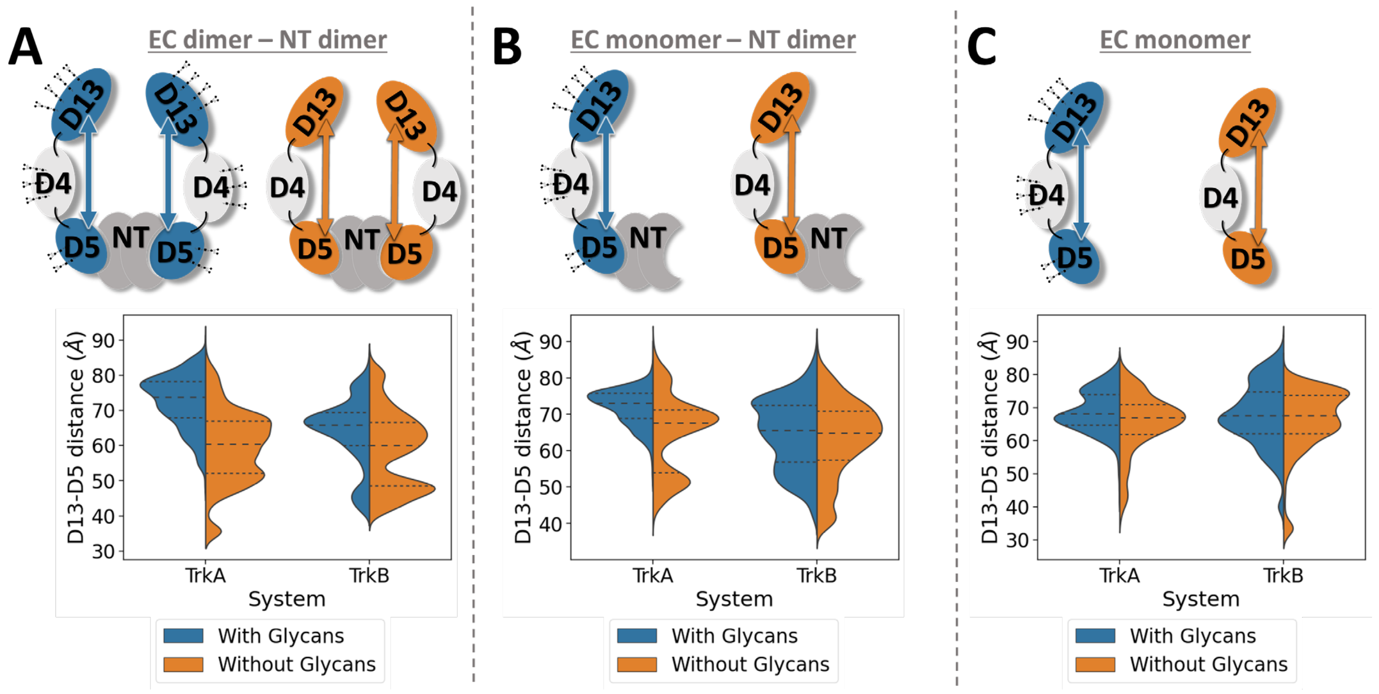


**Figure S26:** Distance between the centers of geometry of the D13-D5 domains in the TrkA and TrkB systems. The distances were calculated every 100 frames and the final distributions for the glycosylated and non-glycosylated TrkA and TrkB systems are shown in violin plots. The quartiles of the distributions are indicated with dashed lines.

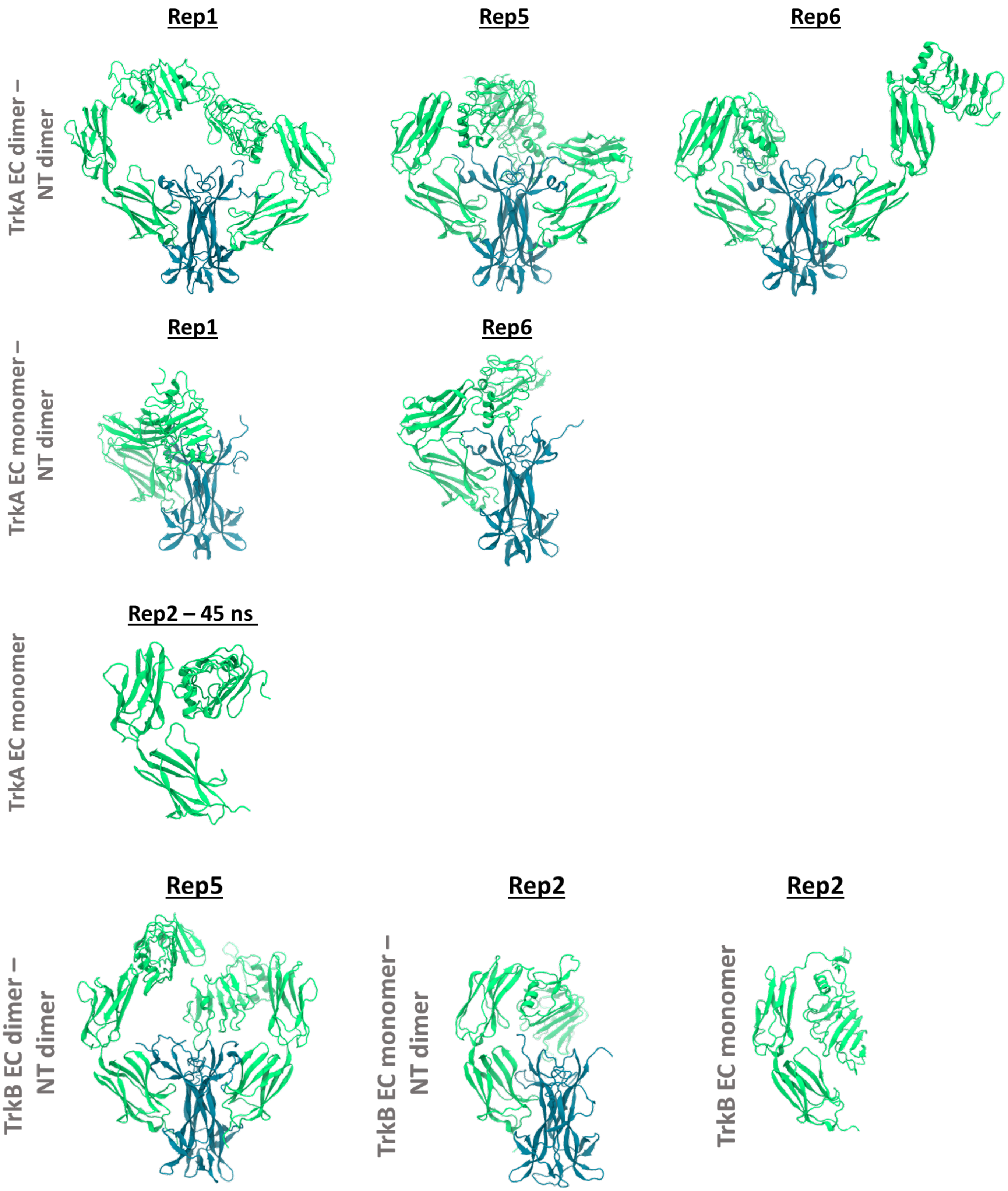


**Figure S27: :** Simulation snapshots of the different non-glycosylated TrkA and TrkB systems that have low values of the radius of gyration and inter-domain distance. All the snapshots are taken from the last frame of the indicated replica simulations, except for the conformation of the TrkA monomer without NT, which is a snapshot at the 45 ns timeframe of the simulation.

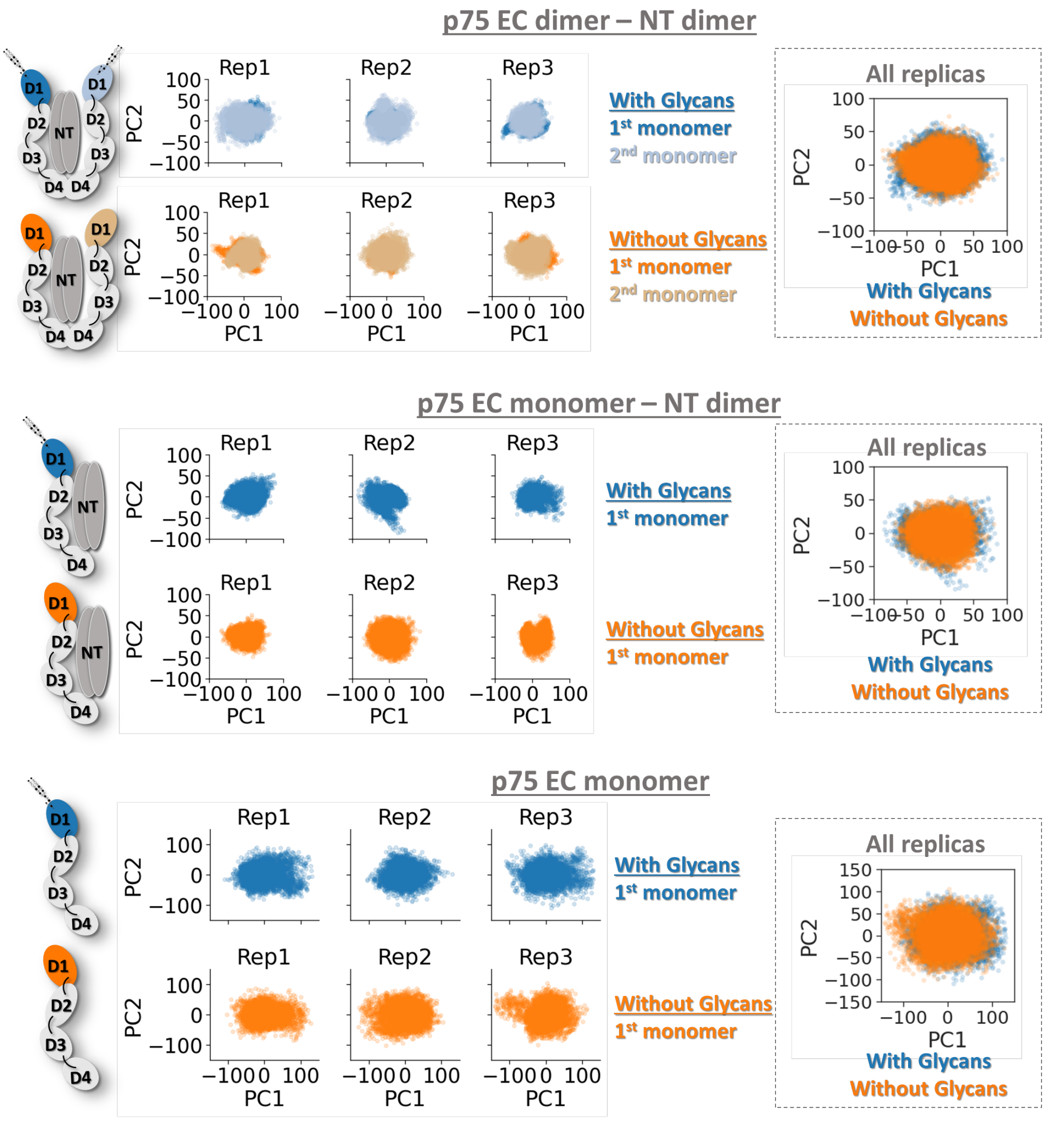


**Figure S28:** Principal component analysis (PCA) of the D1 domain of the p75-ECD after alignment of the structure to the rest of the domains for the glycosylated (blue) and non-glycosylated (orange) systems. The color coding of each domain is shown in the schematic representations of the systems on the left of the plots. Only the 1st and 2nd principal components are displayed.

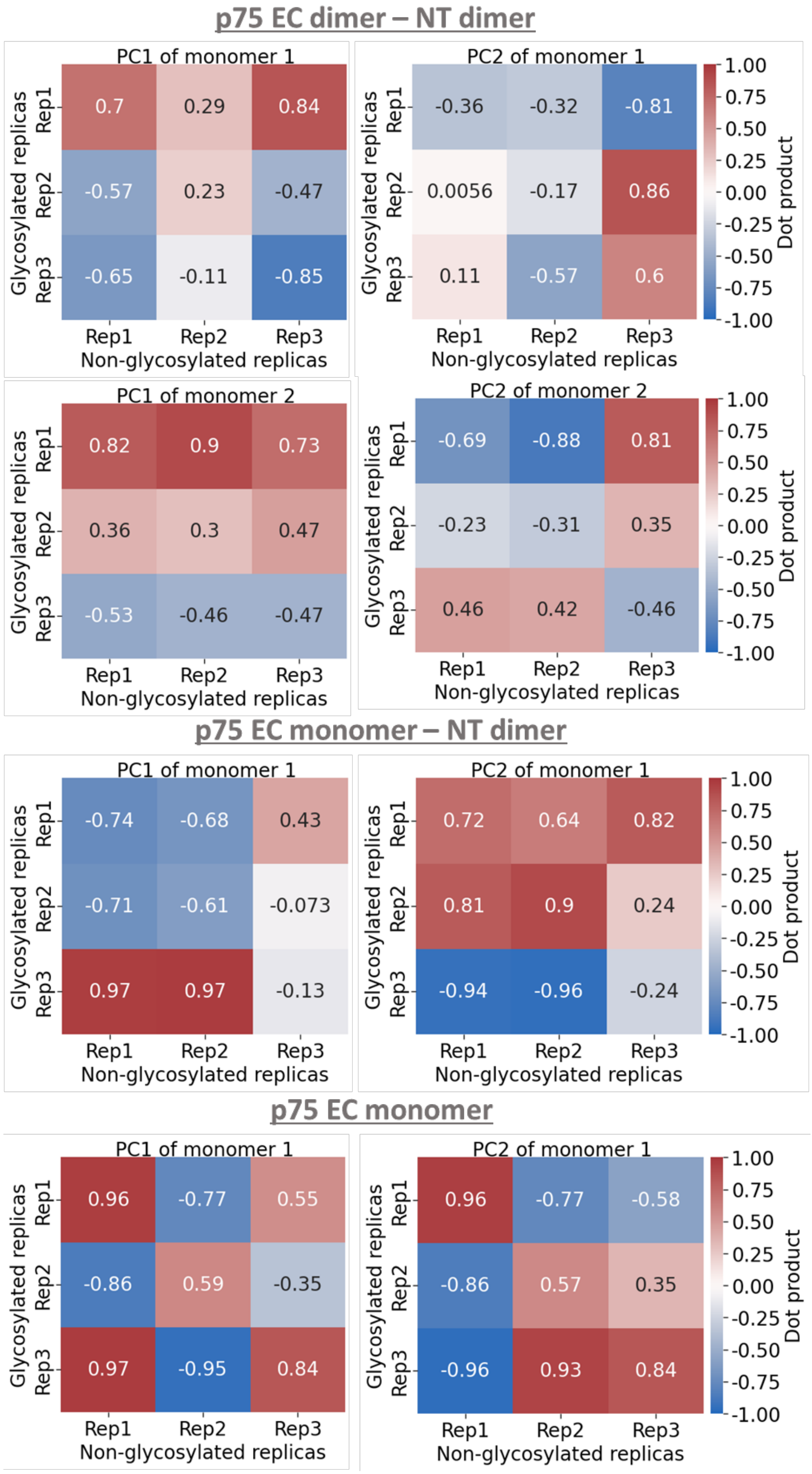


**Figure S29:** Heat maps of the dot products of the 1st and 2nd principal components from the various replicas and systems of the D1 domain of the p75 ECD with and without glycans. Note that PC1 and PC2 do not necessarily correspond to the same vector in all replica simulations. Thus, to compare the PCs between different replicas, the dot products between the PC1s and PC2s from the various replicas were calculated. The results showed that in some cases the PCs were pointing towards the same direction (positive dot products) between the glycosylated and non-glycosylated systems, while in other cases they pointed in opposite directions (negative dot products), or were even perpendicular to each other (close to zero dot products).

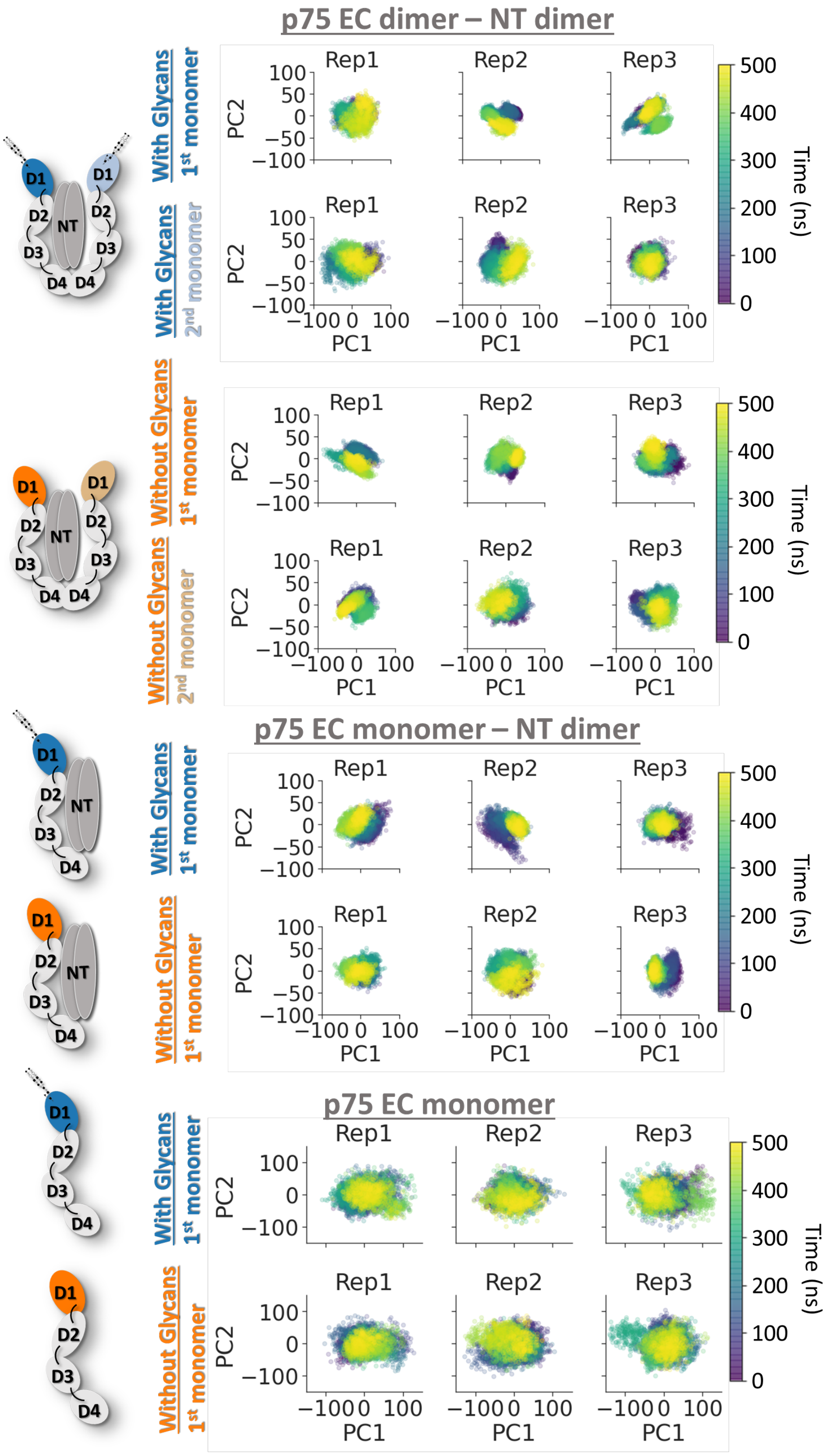


**Figure S30:** Time evolution of the principal components of the D1 domain of the p75-ECD after alignment to the rest of the domains, for the glycosylated and non-glycosylated systems. Only the 1st and 2nd principal components are displayed. During the simulation time, the systems visit the same PC1-PC2 space, indicating the narrow conformational space explored by the system due to the rigidity of the p75-ECD.

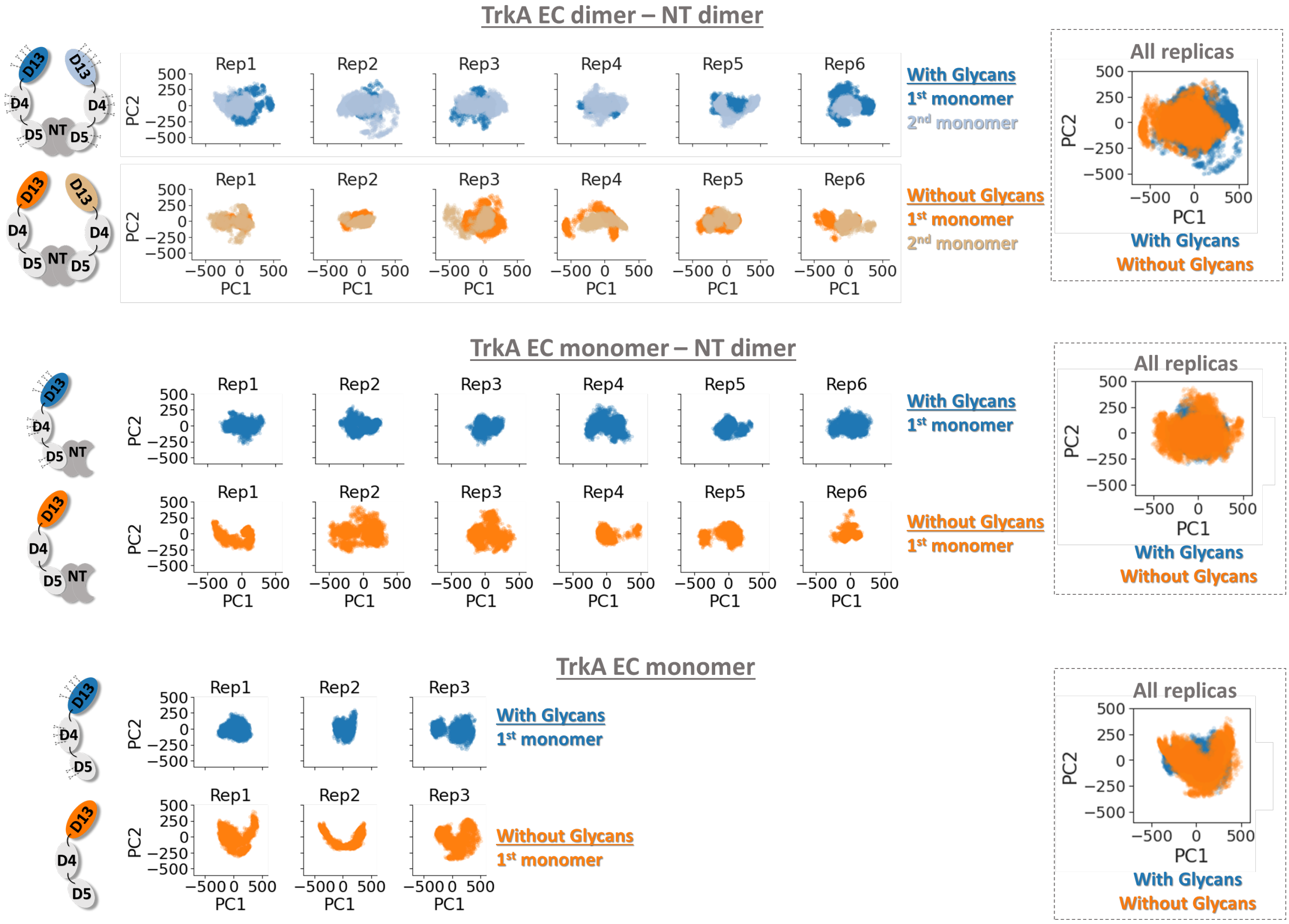


**Figure S31:** Principal component analysis of the D13 domain motions during the simulations of the glycosylated and nonglycosylated TrkA systems. PCA analysis was performed for the D13 domains after alignment of each EC segment to the D4 and D5 domains. The color coding of each domain is shown in the schematic representations of the systems on the left of the plots. Only the 1st and 2nd principal components are displayed.

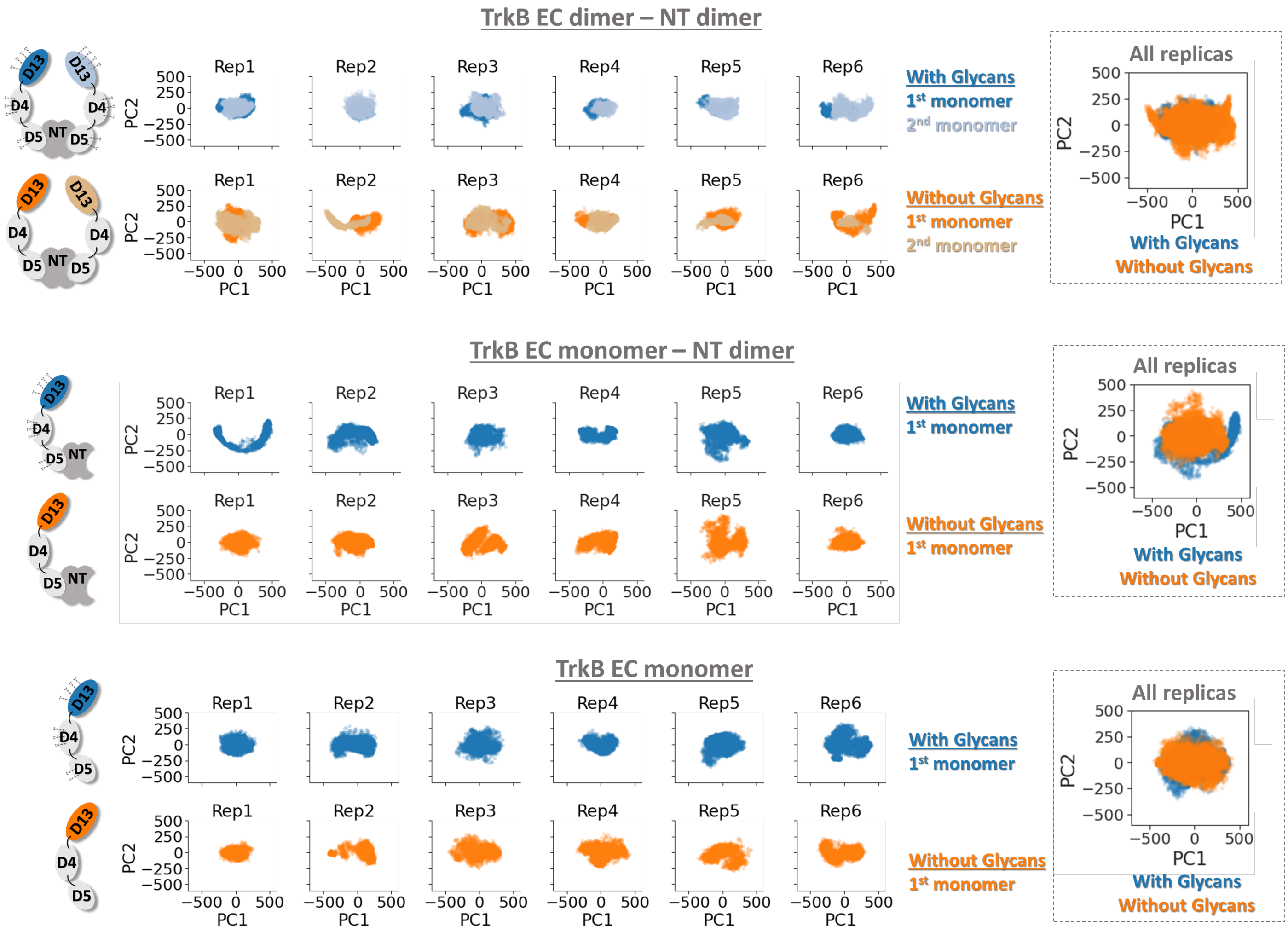


**Figure S32:** Principal component analysis of the D13 domain motions during the simulations of the glycosylated and non-glycosylated TrkB systems. PCA analysis was performed for the D13 domains after alignment of each EC segment to the D4 and D5 domains. The color coding of each domain is shown in the schematic representations of the systems on the left of the plots. Only the 1st and 2nd principal components are displayed.

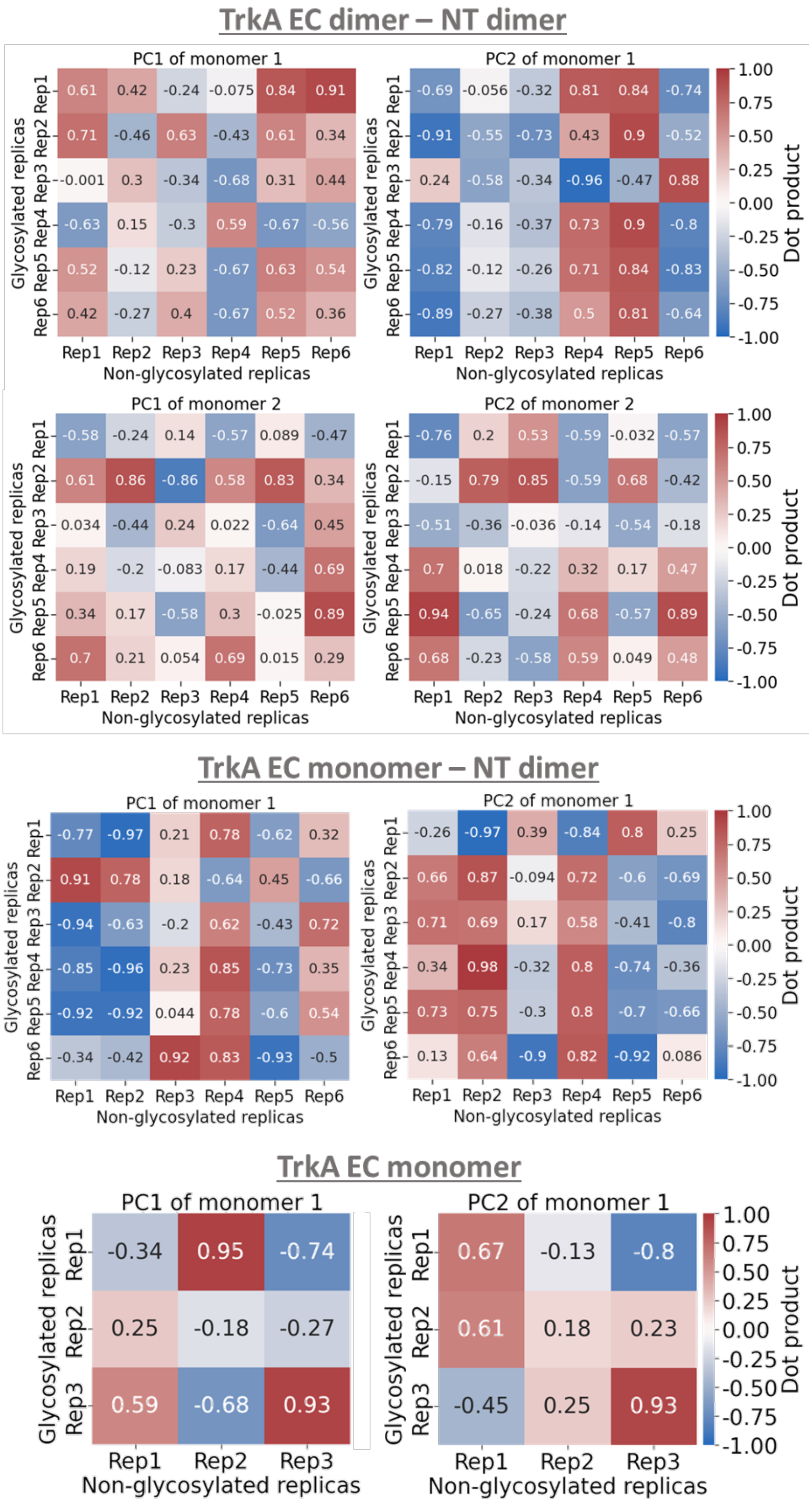


**Figure S33:** Heat maps of the dot products of the first (PC1) and second (PC2) principal components between glycosylated and non-glycosylated replica pairs of the TrkA systems. The PCA was performed for the D13 domain only, after alignment of the trajectories on the D4-D5 domains. In the systems of the receptor dimers, the PC1 and PC2 dot products were calculated for each monomer separately. The dot product values are color-coded, with values equal to 1 (-1) corresponding to parallel (anti-parallel) vectors, and values of 0 indicating perpendicular vectors. The maps indicate a wide distribution of PC vectors, indicating wide exploration of conformation space.

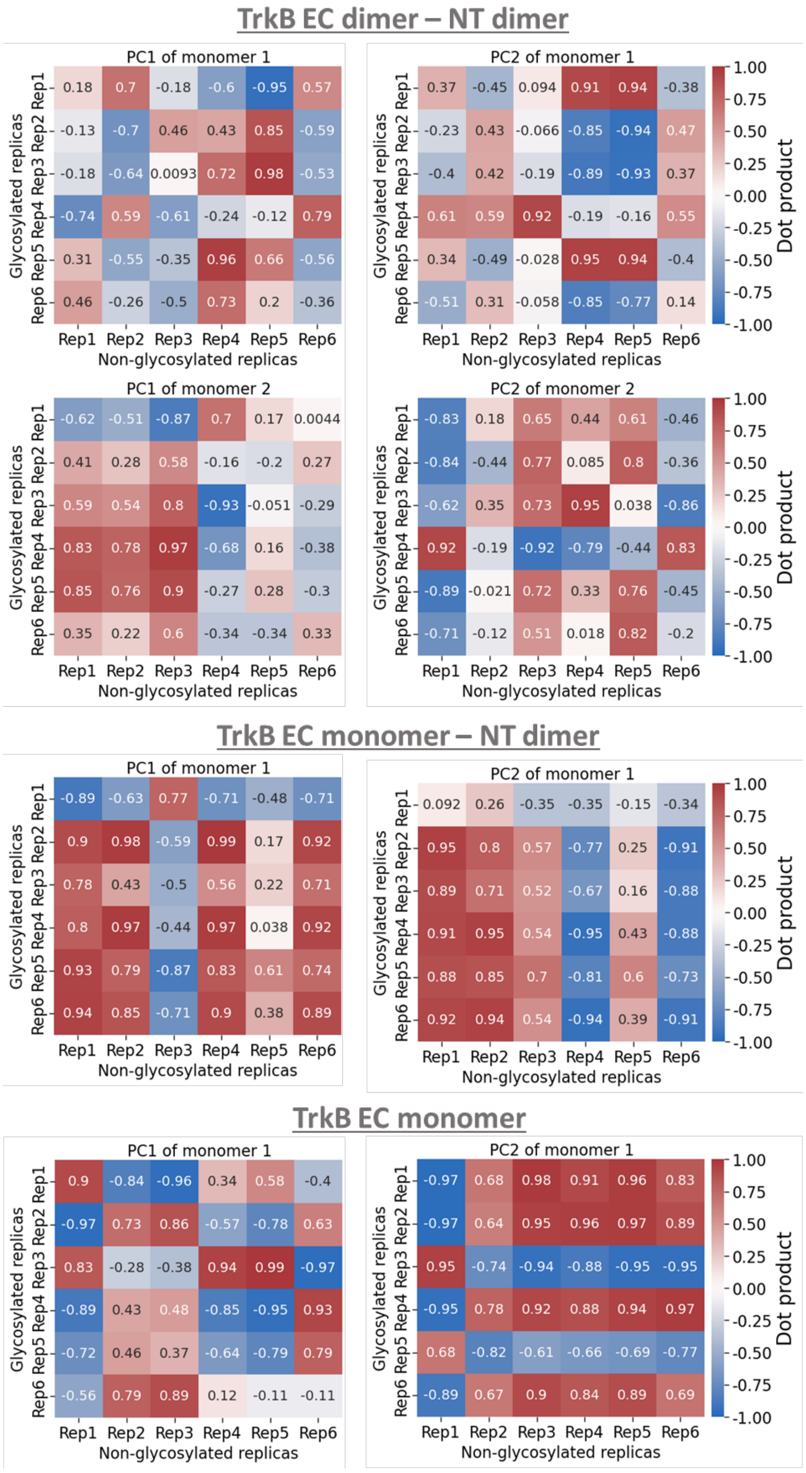


**Figure S34:** Heat maps of the dot products of the first (PC1) and second (PC2) principal components between glycosylated and non-glycosylated replica pairs of the TrkB systems. The PCA was performed for the D13 domain only, after alignment of the trajectories on the D4-D5 domains. In the systems of the receptor dimers, the PC1 and PC2 dot products were calculated for each monomer separately. The dot product values are color-coded, with values equal to 1 (-1) corresponding to parallel (anti-parallel) vectors, and values of 0 indicating perpendicular vectors. In contrast to TrkA, for TrkB, stronger similarities were observed, with parallel or anti-parallel PCs, especially for the systems with receptor monomers. This behavior seems to coincide with the greater differences observed between glycosylated and non-glycosylated forms of the TrkA systems compared to TrkB.

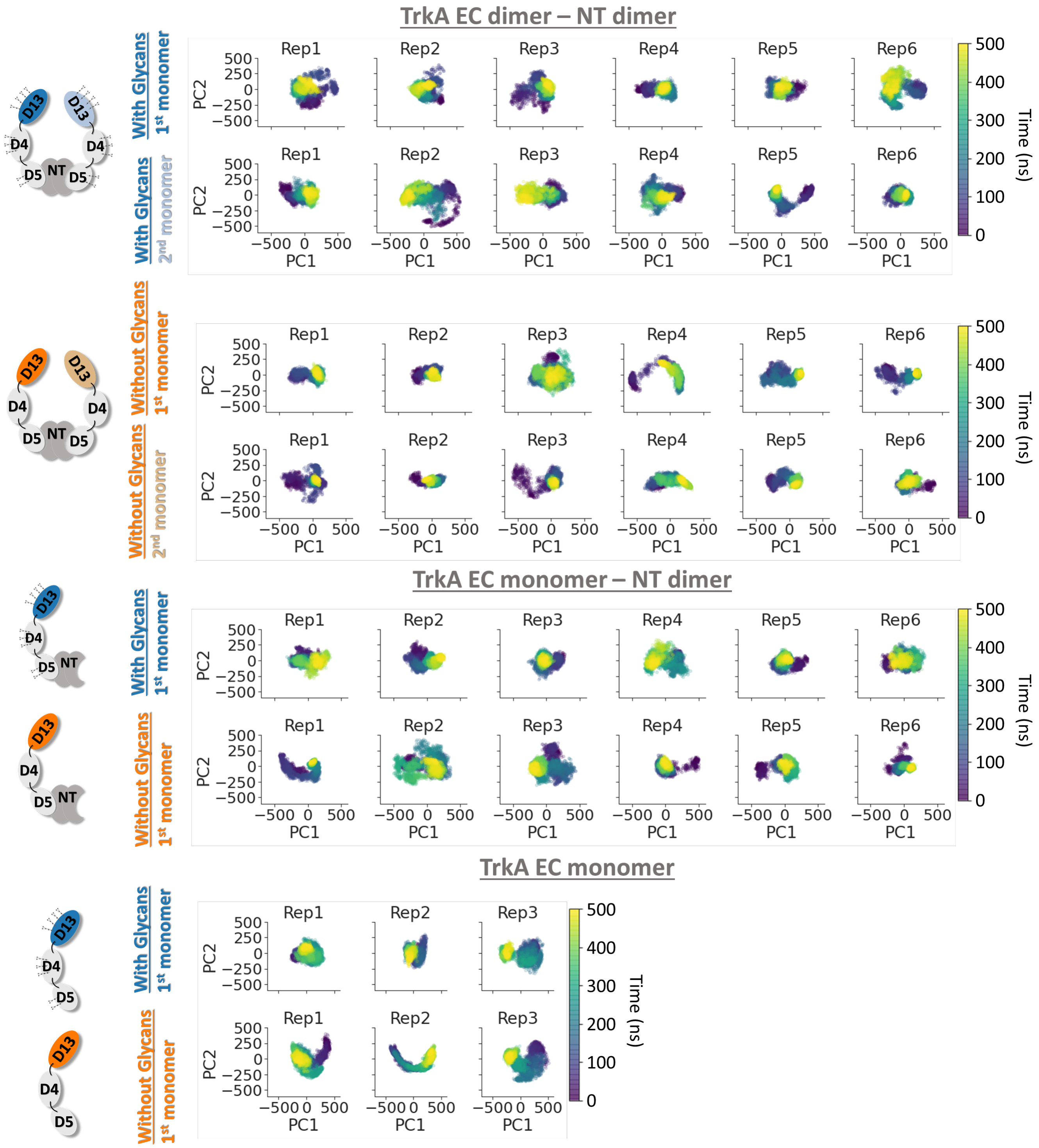


**Figure S35:** Time evolution of the first and second principal components of the D13 domain during the simulations of the glycosylated and not TrkA systems. PCA analysis was performed to the D13 domains after alignment of each EC segment to the D4 and D5 domains. The color coding of each domain is shown in the schematic representations of the systems on the left of the plots. The systems explore specific regions of the PC1 and PC2 space. In cases where conformational change took place during the simulations as indicated by the system moving from one region to another in the PC1/PC2 space, no revisiting of the initial region took place.

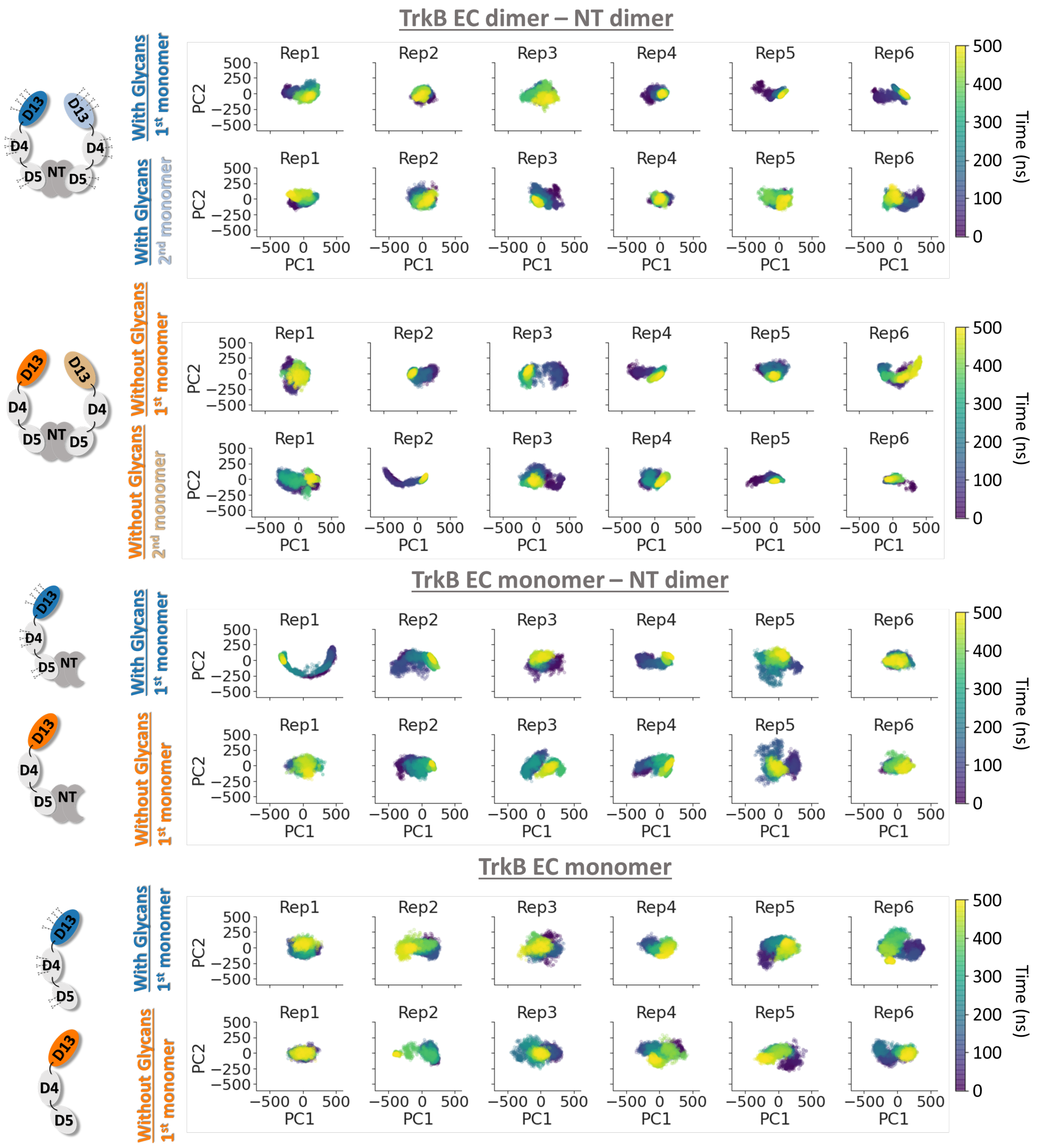


**Figure S36:** Time evolution of the first and second principal components of the D13 domain during the simulations of the glycosylated and not TrkB systems. PCA analysis was performed to the D13 domains after alignment of each EC segment to the D4 and D5 domains. The color coding of each domain is shown in the schematic representations of the systems on the left of the plots. In cases where conformational change took place during the simulations as indicated by the system moving from one region to another in the PC1/PC2 space, no revisiting of the initial region took place.

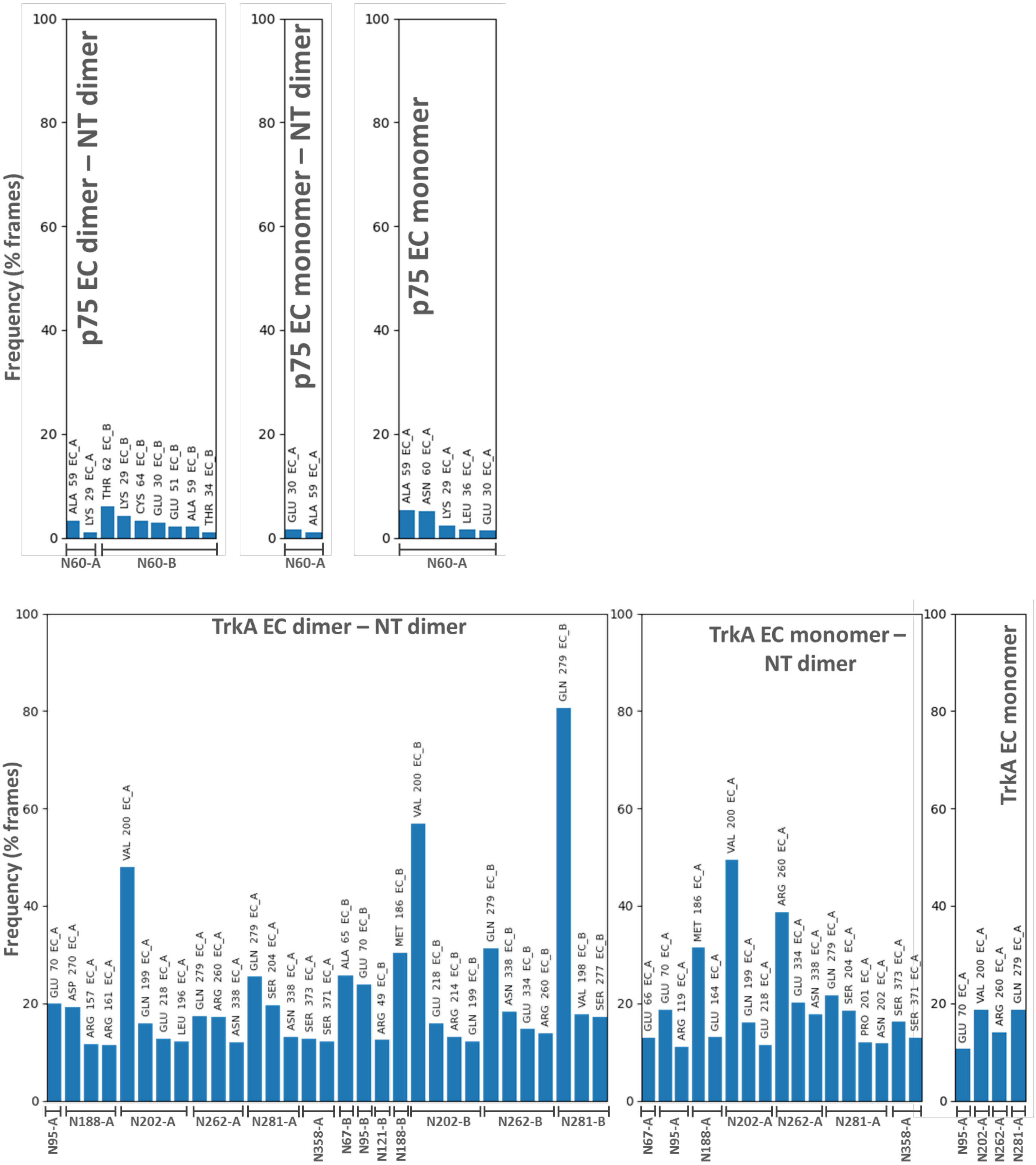
p75 EC dimer – NT dimer
p75 EC monomer – NT dimer
p75 EC monomer
Frequency (% frames)
ALA 59 EC_A
LYS 29 EC_A
THR 62 EC_B
LYS 29 EC_B
CYS 64 EC_B
GLU 30 EC_B
GLU 51 EC_B
ALA 59 EC_B
THR 34 EC_B
N60-A
N60-B
GLU 30 EC_A
ALA 59 EC_A
N60-A
ALA 59 EC_A
ASN 60 EC_A
LYS 29 EC_A
LEU 36 EC_A
GLU 30 EC_A
N60-A
TrkA EC dimer – NT dimer
TrkA EC monomer – NT dimer
TrkA EC monomer
Frequency (% frames)
GLU 70 EC_A
ASP 270 EC_A
ARG 157 EC_A
ARG 161 EC_A
VAL 200 EC_A
GLN 199 EC_A
GLU 218 EC_A
LEU 196 EC_A
GLN 279 EC_A
ARG 260 EC_A
ASN 338 EC_A
GLN 279 EC_A
SER 204 EC_A
ASN 338 EC_A
SER 373 EC_A
SER 371 EC_A
ALA 65 EC_B
GLU 70 EC_B
ARG 49 EC_B
MET 186 EC_B
VAL 200 EC_B
GLU 218 EC_B
ARG 214 EC_B
GLN 199 EC_B
GLN 279 EC_B
ASN 338 EC_B
GLU 334 EC_B
ARG 260 EC_B
GLN 279 EC_B
VAL 198 EC_B
SER 277 EC_B
N95-A
N188-A
N202-A
N262-A
N281-A
N358-A
N67-B
N95-B
N121-B
N188-B
N202-B
N262-B
N281-B
GLU 66 EC_A
GLU 70 EC_A
ARG 119 EC_A
MET 186 EC_A
GLU 164 EC_A
VAL 200 EC_A
GLN 199 EC_A
GLU 218 EC_A
ARG 260 EC_A
GLU 334 EC_A
ASN 338 EC_A
GLN 279 EC_A
SER 204 EC_A
PRO 201 EC_A
ASN 202 EC_A
SER 373 EC_A
SER 371 EC_A
N67-A
N95-A
N188-A
N202-A
N262-A
N281-A
N358-A
GLU 70 EC_A
VAL 200 EC_A
ARG 260 EC_A
GLN 279 EC_A
N95-A
N202-A
N262-A
N281-A

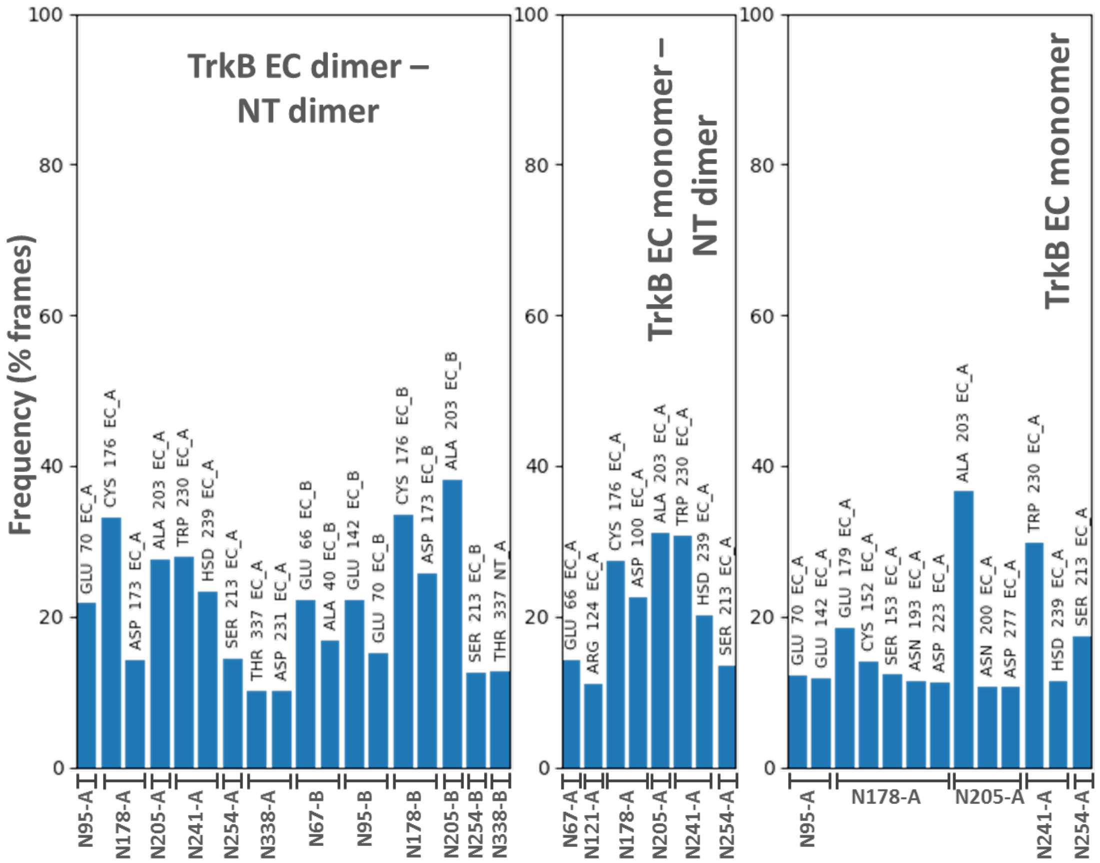


**Figure S37:** Contact analysis between the glycans attached to different Asn residues and the protein residues in all glycosylated p75, TrkA and TrkB systems. A contact is defined when any non-hydrogen atom of a glycan is within 4.5 Å distance of any non-hydrogen atom of a protein residue. The glycans are denoted by the Asn residues of each receptor monomer (A or B) on the x-axis, while the protein residues that interact with the corresponding glycan are indicated on top of each bar. The height a bar shows the percentage of the total simulation time in which the corresponding contact is present. Only contacts with occupancy higher than 1% for p75, and 10% for the Trks are shown.
